# Y-complex nucleoporins independently contribute to nuclear pore assembly and gene regulation in neuronal progenitors

**DOI:** 10.1101/2023.01.24.524209

**Authors:** Clarisse Orniacki, Annalisa Verrico, Stéphane Pelletier, Benoit Souquet, Fanny Coulpier, Laurent Jourdren, Serena Benetti, Valérie Doye

## Abstract

From their essential function in building up the nuclear pore complexes, nucleoporins have expanded roles beyond nuclear transport. Hence, their contribution to chromatin organization and gene expression has set them as critical players in development and pathologies. We previously reported that Nup133 and Seh1, two components of the Y-complex subunit of the nuclear pore scaffold, are dispensable for mouse embryonic stem cell viability but required for their survival during neuroectodermal differentiation. Here, a transcriptomic analysis revealed that Nup133 regulates a subset of genes at early stages of neuroectodermal differentiation, including *Lhx1 and Nup210L*, encoding a newly validated nucleoporin. These genes were also misregulated in *Nup133∆Mid* neuronal progenitors, in which NPC basket assembly is impaired, as previously observed in pluripotent cells. However, a four-fold reduction of Nup133, despite affecting basket assembly, is not sufficient to alter *Nup210L* and *Lhx1* regulation. Finally, these two genes are also misregulated in *Seh1*-deficient neural progenitors that only show a mild decrease in NPC density. Together these data reveal a shared function of Y-complex nucleoporins in gene regulation during neuroectodermal differentiation, which seem independent of nuclear pore basket assembly.

## INTRODUCTION

As channels embedded in the nuclear envelope, the nuclear pore complexes (NPCs) constitute the only gateway for selective transport of macromolecules between the cytoplasm and the nucleus. These impressive structures are composed of proteins called nucleoporins (Nups) that assemble in a highly organized and modular manner (reviewed in Dultz et al., (2022)). The Y-complex - also named Nup107-160 complex - that comprises in vertebrates nine distinct proteins, is a key structural subunit of the NPC scaffold. 16 copies of this complex assemble on the nuclear and cytoplasmic sides of the NPC to build up its outer rings, to which cytoplasmic filaments and the nuclear basket are anchored.

In addition to their canonical nuclear transport function, many Nups are also known to have other cellular functions, notably in cell cycle progression or as key regulators of chromatin organization and gene expression (reviewed in Buchwalter et al., 2019; Hezwani and Fahrenkrog, 2017). In line with these multiple functions, mutations in many Nups have been identified as primary causes of rare genetic diseases. Despite the presence of NPCs in all nucleated cells, most of these diseases specifically affect one or a few organs (reviewed in Jühlen and Fahrenkrog, 2018). Such tissue or cell-type specific alterations may reflect variable Nup stoichiometry at NPCs, as notably reported for several integral membrane Nups and peripheral Nups (Ori et al., 2013). For instance, increased levels of the transmembrane protein Nup210 in myoblasts and neuronal progenitors was shown to be critical for their differentiation (D’Angelo et al., 2012). Likewise, depletion of the basket nucleoporin Nup50 reduces the differentiation efficiency of C2C12 myoblasts (Buchwalter et al., 2014). In contrast, another basket nucleoporin, Nup153, which is highly expressed in pluripotent cells and neuronal progenitors compared to differentiated cells, is required for the maintenance of their identities, notably by regulating epigenetic gene silencing and transcriptional programs (Jacinto et al., 2015; Toda et al., 2017). More recently, the Y-complex constituent Seh1, which is highly expressed in oligodendrocyte progenitor cells, was shown to be required for oligodendrocyte differentiation and myelination by regulating the assembly of a transcription complex at the nuclear periphery (Liu et al., 2019). However, individual Y-complex Nups also contribute to earlier stages of differentiation, as underscored by the impaired neuroectodermal differentiation of *Nup133*^*-/-*^, *Seh1*^*-/-*^ and *Nup43*^*-/-*^ mouse embryonic stem cells (mESCs) (Gonzalez-Estevez, Verrico et al., 2021; Lupu et al., 2008).

Earlier studies had found that the vertebrate Y-complex is, as an entity, critically required for NPC assembly both at the end of mitosis and during interphase (Doucet and Hetzer, 2010; Harel et al., 2003; Vollmer et al., 2015; Walther et al., 2003). The viability of *Nup133*^*-/-*^, *Seh1*^*-/-*^ and *Nup43*^*-/-*^ mESCs however indicated that the corresponding Y-complex Nups were individually largely dispensable for nuclear pore assembly in these pluripotent cells. Consistently, we previously showed that mutations of Nups that form the short arm of the Y-complex, namely Nup43, Nup85 and Seh1, only lead to a mild decrease in NPC density in pluripotent mESCs (Gonzalez-Estevez, Verrico et al., 2021). In contrast, pluripotent *Nup133*^*-/-*^ mESCs feature a normal NPC density, but show specific nuclear basket defects, with half of NPCs lacking Tpr while Nup153 dynamics was increased (Souquet et al., 2018). How Y-complex Nups contribute to NPC assembly in differentiating mESCs was unknown.

Because of the established implication of the basket nucleoporins Nup153 and Tpr in chromatin organization and gene regulation (Aksenova et al., 2020; Boumendil et al., 2019; Jacinto et al., 2015; Krull et al., 2010; Toda et al., 2017), we decided to investigate potential gene expression defects in *Nup133*^*-/-*^ mESCs during neuroectodermal differentiation. Here we show that Nup133 regulates a subset of genes, including *Lhx1* and *Nup210L* that are similarly misregulated in the absence of Seh1. However, *Nup133* and *Seh1* deficiencies display distinct NPC assembly phenotypes in neuronal progenitors, thus indicating separate roles for these proteins in NPC architecture and gene regulation in the context of mESC differentiation.

## RESULTS AND DISCUSSION

### Nup133 is required for the regulation of a subset of genes during neuroectodermal differentiation

The impaired neuroectodermal differentiation of *Nup133*^*-/-*^ mESCs initially described in Lupu et al. (2008), was also observed in HM1-derived *Nup133*^*-/-*^mESCs as revealed by their altered growth and increased cell death (***Figure 1A, B***). To assess the potential effect of Nup133 deficiency in gene regulation upon neuroectodermal differentiation, we first determined the mRNA levels of genes expressed in pluripotent cells (*Oct4* and *Nanog*) and in early neuronal progenitors (*Sox1* and *Pax6*) that are considered markers for the respective states. RT-qPCR analyses showed that these genes were properly repressed and activated, respectively, in *Nup133*^*-/-*^ cells stimulated to differentiate towards neuroectoderm (***Figure 1C***). This indicated that despite their impaired viability at early stages of differentiation (***Figure 1A, B***) the surviving *Nup133*^*-/-*^ cells are able to exit pluripotency and to commit towards the neuronal lineage, without overt defects in the expression of these markers.

**Figure 1.**
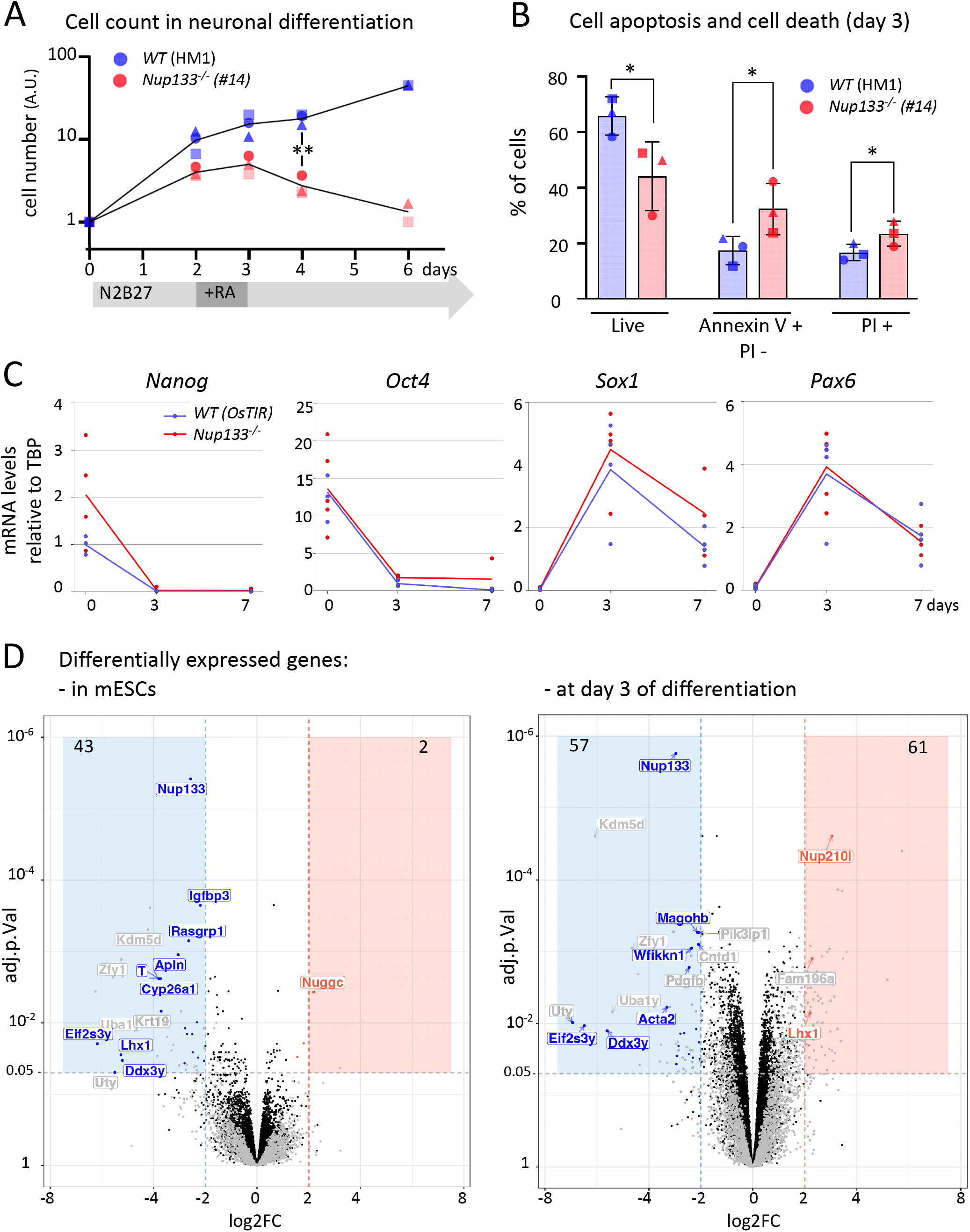
Transcriptomic analysis of *Nup133*^*-/-*^ mESCs and early neuronal progenitors. **A**. Growth curve in neuroectodermal differentiation of *WT* (HM1) and isogenic *Nup133*^*-/-*^ mESCs. The graph (logarithmic scale) represents the average of cell counts from 3 independent experiments, each represented by a distinct label. Values were normalized to the number of cells seeded at day 0 (3.10^4^ cells/cm^2^). **B**. Quantification of apoptosis initiation (defined by annexin V positive (+) and propidium iodide negative (PI -) cells) and cell death (propidium iodide positive cells, PI+) in *WT* and *Nup133*^*-/-*^ cells at day 3 of differentiation. Error bars represent standard deviation of 3 independent experiments, each represented by a distinct label. In A and B, significance was assessed between *WT* and *Nup133*^*-/-*^ using paired T. test (*p<0.5; **p<0.01). **C**. mRNA levels of pluripotency (*Nanog, Oct4*) and neuronal progenitor (*Sox1, Pax6*) markers during neuroectodermal differentiation were quantified by RT-qPCR and normalized to *Tbp* mRNA levels. Each dot represents an individual experiment. **D**. Volcano plots of the RNA-seq analysis carried out in pluripotent mESCs and in cells at day 3 of neuroectodermal differentiation, showing differentially expressed genes (DEGs), by fold change (log2FC of *Nup133*^*-/-*^ compared to *WT* cells) and significance (adj. p.Val presented on a -log10 scale). Significantly upregulated DEGs (adj. p-value<0.05, logFC>2) and downregulated DEGs (adj. p-value<0.05, logFC<-2) are represented by red and blue dots, respectively, if their average normalized expression in log2(CPM) is above 1. Their number is indicated at the top of each colored square. Among them, the names of DEGs assessed by RT-qPCR are indicated in blue or red. The names of other relevant DEGs are indicated in grey. The other genes are represented as grey dots when their average expression is below 1 and otherwise as black dots.

To more broadly explore the impact of Nup133 on gene expression, we compared the transcriptome of *WT* and *Nup133*^*-/-*^ mESCs at the pluripotent state and after 2 or 3 days of differentiation towards neuroectodermal lineage. We therefore used cell lines from two distinct genetic backgrounds, namely the HM1 control cell line and its CRISPR/Cas9-edited *Nup133*^*-/-*^ derivatives (#14 and #19), and the blastocyst-derived control (#1A4) and *Nup133*^*-/-*^ (*merm*, #319) mESC lines (Souquet et al., 2018 and Table S1). This analysis revealed that the transcriptomes of pluripotent *WT* and *Nup133*^*-/-*^ mESCs were overall similar, whereas an increasing number of genes were misregulated at day 3 of differentiation (***Figures 1D and S1A***).

We assayed by RT-qPCR the altered expression of a subset of these genes, filtered by criteria of differential expression (logFC>2 or <-2), significance (adjusted p-value<0.05) and lastly by average expression level (average number of reads with a log2(CPM)>1, to ensure proper detection by RT-qPCR) (***Figure S1B-E***). In addition to *WT (HM1)* and *Nup133*^*-/-*^ (#14) used for the initial RNA-seq experiment, these analyses were also conducted on samples from *Nup133* “*Rescue*” cell lines generated by inserting the GFP-Nup133 transgene in *Nup133*^*-/-*^ (#14) mESCs at the permissive *Tigre* locus (Zeng et al., 2008). As an additional control, we used an HM1-derived cell line that carries a transgene (*OsTIR*) similarly inserted in the *Tigre* locus (***Figure S2A and Table S1***). In contrast to the impaired viability of *Nup133*^*-/-*^ mESCs upon neuroectodermal differentiation, the survival of the *Rescue* and *WT (OsTIR)* cell lines were similar, confirming the functionality of the GFP-Nup133 transgene (***Figure 2A-C)***.

**Figure 2.**
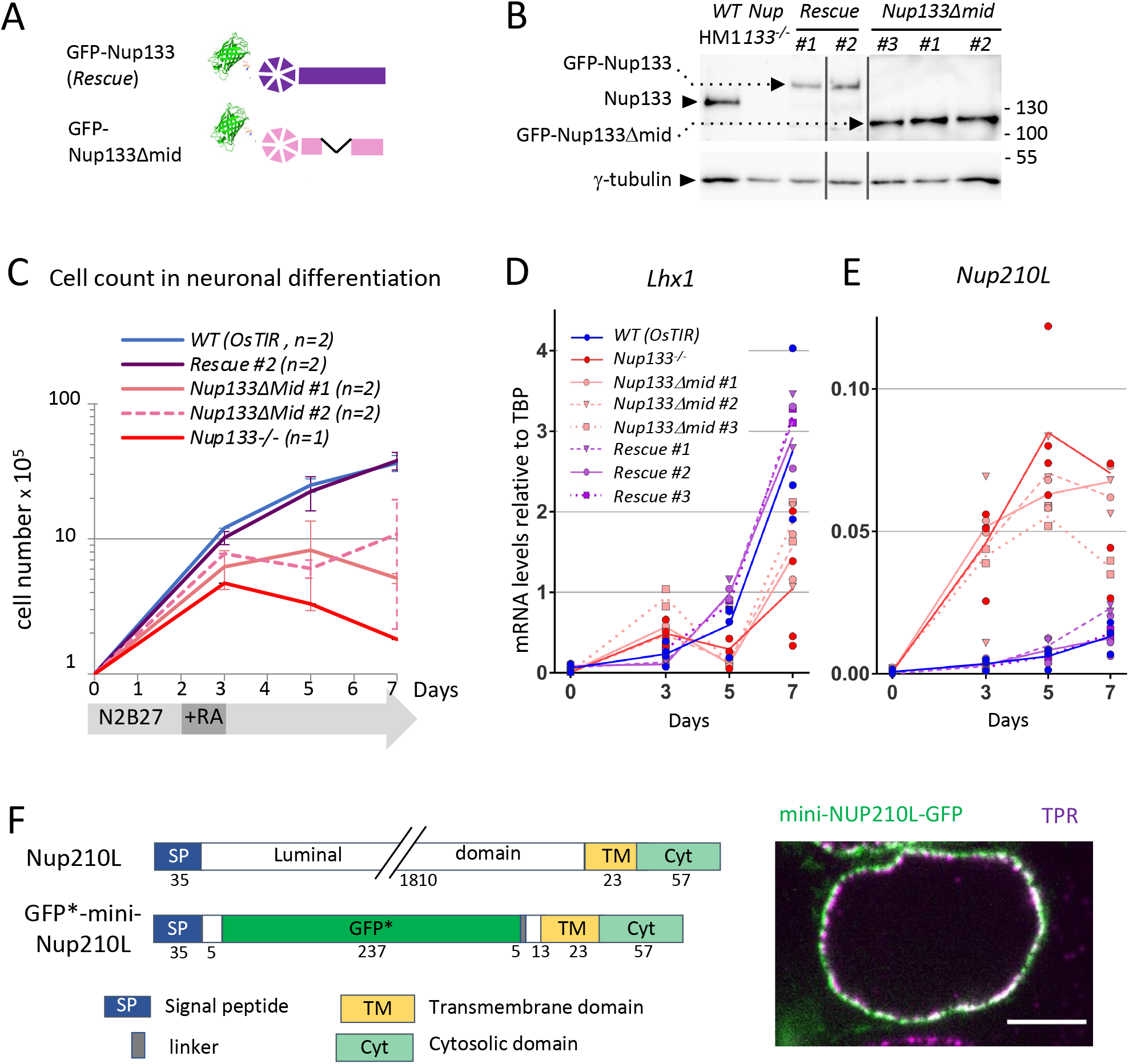
Altered expression of *Lhx1* and *Nup210L* in *Nup133* mutant cell lines. **A**. Schematics of the GFP-Nup133 (*Rescue*) and GFP-Nup133∆mid fusion proteins (see also Table S1 and Figure S2A). **B**. Western blot showing the expression of endogenous Nup133 or its GFP-tagged forms in *WT, Nup133*^*-/-*^, *Nup133-Rescue* and *Nup133∆mid* cells at day 7 of differentiation. γ-tubulin is used as loading control. Molecular weights are indicated (kilodaltons). **C**. Growth curve obtained from cell counts in neuroectodermal differentiation (two independent experiments) presented in a logarithmic scale. Cells were seeded at a density of ∼3.10^4^ cells/cm^2^ (10^5^ cells in p12 wells). Error bars correspond to standard deviations. **D, E**. *Lhx1* and *Nup210L* mRNA levels were analyzed by RT-qPCR at the indicated time points of neuroectodermal differentiation in *WT (OsTIR), Nup133*^*-/-*^, *Rescue* and *Nup133∆mid* cell lines. Each cell line is represented by a distinct label. **F**. Left: Schematics of Nup210L and of the GFP*-mini-Nup210L construct that comprises: Nup210L predicted signal peptide (SP) and part of the luminal domain [aa 1-40], GFP* (see details about GFP* in the legend to Figure S2) and aa 1791-1884 of Nup210L, encompassing part of its predicted luminal domain and its transmembrane TM and cytosolic (Cyt) domains. Numbers below the schematics correspond to amino acid residues. Right: *WT* (*HM1)* mESCs transiently expressing the GFP*-mini-Nup210L construct were fixed and processed for immunofluorescence with anti-Tpr antibodies. A single confocal section is shown. Scale bar, 5µm.

For the candidates genes localized on the short arm of the Y chromosome (*Ddx3y* and *Eif2s3y*, also located in close proximity to the loci of *Uty, Uba1y, Kdm5d and Zfy* (Subrini and Turner, 2021)) we observed clone-dependent expression variations (***Figure S1B)***. This suggests that their apparent shared misregulations, also reported in *Tet1* and *Tet2* mutant mESCs (Huang et al., 2013), might be in our case caused by partial loss or silencing of this genomic region, independently of *Nup133* deficiency.

In contrast, we could validate the increased mRNA levels *in Nup133*^*-/-*^ compared to *WT* of *Nuggc* at day 0, and of *Nup210L* and *Lhx1* at day 3 of differentiation (***Figure S1C***). We also confirmed the reduced mRNA levels in *Nup133*^*-/-*^ compared to *WT* for *Magohb* and *Wfikkn1* (but not *Acta2*) at day 3 of differentiation (***Figure S1D***). Finally, reduced mRNA levels of the assayed candidate genes at the pluripotent state (day 0) were not significant due to high variability among replicates or between control cell lines (HM1 and OsTIR) (***Figure S1E***). Importantly, among the validated candidate genes, *Lhx1, Nup210L, Nuggc* and *Magohb* were all efficiently restored to wild-type levels by the *GFP-Nup133* transgene.

*Lhx1* is a transcription factor involved in kidney and brain differentiation (Costello et al., 2015; Delay et al., 2018; McMahon et al., 2019; Shawlot et al., 1999), two organs affected in rare genetic diseases linked to *Nup133* mutations, namely steroid-resistant nephrotic syndrome and Galloway Mowat syndrome (Braun et al., 2018; Fujita et al., 2018). We further focused on this gene because of its complex misregulation in *Nup133*^*-/-*^ cells. Indeed, while we confirmed by RT-qPCR the upregulation of *Lhx1* expression at day 3 of differentiation (***Figure S1C***), *Lhx1* was subsequently downregulated again at later time-points (days 5 and 7 of differentiation) in *Nup133*^*-/-*^ cells compared to *WT* and *Rescue* cells (***Figure 2D***).

The other candidate gene we further characterized, *Nup210L*, is the differentially expressed gene (DEG) with the most significant p-value at day 3 of differentiation (***Figure 1D***). It is also one of the rare DEGs whose expression already increased at day 2 compared to *WT* cells (***Figure S1A***). In mice, *Nup210L* mRNA is mainly detected in the testis and to a lesser extent in the embryonic brain (https://www.ncbi.nlm.nih.gov/gene/77595); in humans, besides testis, *Nup210L* expression was also detected in the prefrontal cortex neurons of rare individuals (Gusev et al., 2019). Analyses at later stages (day 5 and 7) of differentiation towards neuroectoderm showed that *Nup210L* was still more expressed in *Nup133*^*-/-*^ compared to *WT* and *Rescue* cells, although a progressive increase of its expression was also observed in the latter cell lines (***Figure 2E***).

As its name implies, *Nup210L* encodes a potential homologue of the transmembrane nucleoporin Nup210/gp210. However, its putative NPC localization had never been established. To address this issue, we generated a GFP-tagged construct encompassing the minimal NPC targeting determinants previously established for gp210/Nup210 (Wozniak et al., 1989), namely Nup210L predicted signal peptide, transmembrane domain and C-terminal domain (***Figure 2F***). This Nup210L-mini construct when expressed in mESCs colocalizes at the NPC with Tpr. This indicates that, like its homolog, Nup210L is indeed a nucleoporin.

### The middle-domain of Nup133 is required for mESC differentiation, gene regulation and nuclear basket assembly in neuronal progenitors

Having established the requirement of Nup133 for cell viability upon differentiation and for the regulation of a subset of genes, we next aimed to determine how Nup133 contributes to these processes. We first focused on the middle domain of Nup133 that is necessary for the proper assembly of the nuclear pore basket in pluripotent mESCs (Souquet et al., 2018). We therefore integrated the pCAG-GFP-Nup133∆mid transgene in HM1-derived *Nup133*^*-/-*^ mESCs at the *Tigre* locus (***Figure S2A***). The GFP-Nup133∆mid protein levels in the resulting *Nup133∆mid* cells were comparable to those of endogenous Nup133 and of GFP-Nup133 in the *Rescue* cell lines throughout differentiation (***Figure 2B***). Cell counts upon monolayer differentiation towards neuroectodermal lineage showed for the *Nup133∆mid* cell lines a viability phenotype intermediate between *WT* and *Nup133*^*-/-*^, indicating that the middle domain of Nup133 is required for some, but not all of the functions of this nucleoporin upon differentiation (***Figure 2C***).

In contrast, RT-qPCR analysis showed that *Nup210L* and *Lhx1* were similarly misregulated upon neuronal differentiation in *Nup133∆mid* and in *Nup133*^*-/-*^ cells (***Figure 2D and E***). The improved survival upon differentiation of *Nup133∆mid* compared to *Nup133*^*-/-*^ cells enabled us to perform immunofluorescence analyses to determine whether the NPC basket assembly defects, previously observed in pluripotent mESCs lacking Nup133 or its middle domain, also occurred at the differentiated stage. Quantitative immunofluorescence analyses, performed after 5 days of differentiation, showed that the intensity of Tpr at the nuclear envelope was comparable between the *WT* and *Rescue* cell lines. In contrast, a two-fold decrease was observed in *Nup133∆mid* neuronal progenitors (***Figure 3A, B***), a defect comparable to the one previously observed at the pluripotent state (Souquet et al., 2018). In addition, we also measured an increased Nup153 intensity at the nuclear envelope in *Nup133∆mid* progenitors compared to neuronal progenitors from *WT* or *Rescue* cell lines (***Figure 3C***). This increased level of Nup153 is unlikely to solely reflect a global increase in NPC number as reported upon Tpr depletion in other cell lines (McCloskey et al., 2018), since similar quantifications revealed a milder increase of Nup98 intensity at the nuclear envelope compared to Nup153 (***Figure 3D***). Likewise, an increased accessibility of the Nup153 epitope when Tpr is absent seems unlikely, as such an effect was not previously observed in Nup133^-/-^ mESCs at the pluripotent stage (Souquet et al., 2018). The high level of Nup153 observed may therefore reflect an increased stoichiometry of Nup153 per NPC in *Nup133∆mid-* compared to control-derived neuronal progenitors, possibly reflecting different stages of differentiation as previously described (Toda et al., 2017).

**Figure 3.**
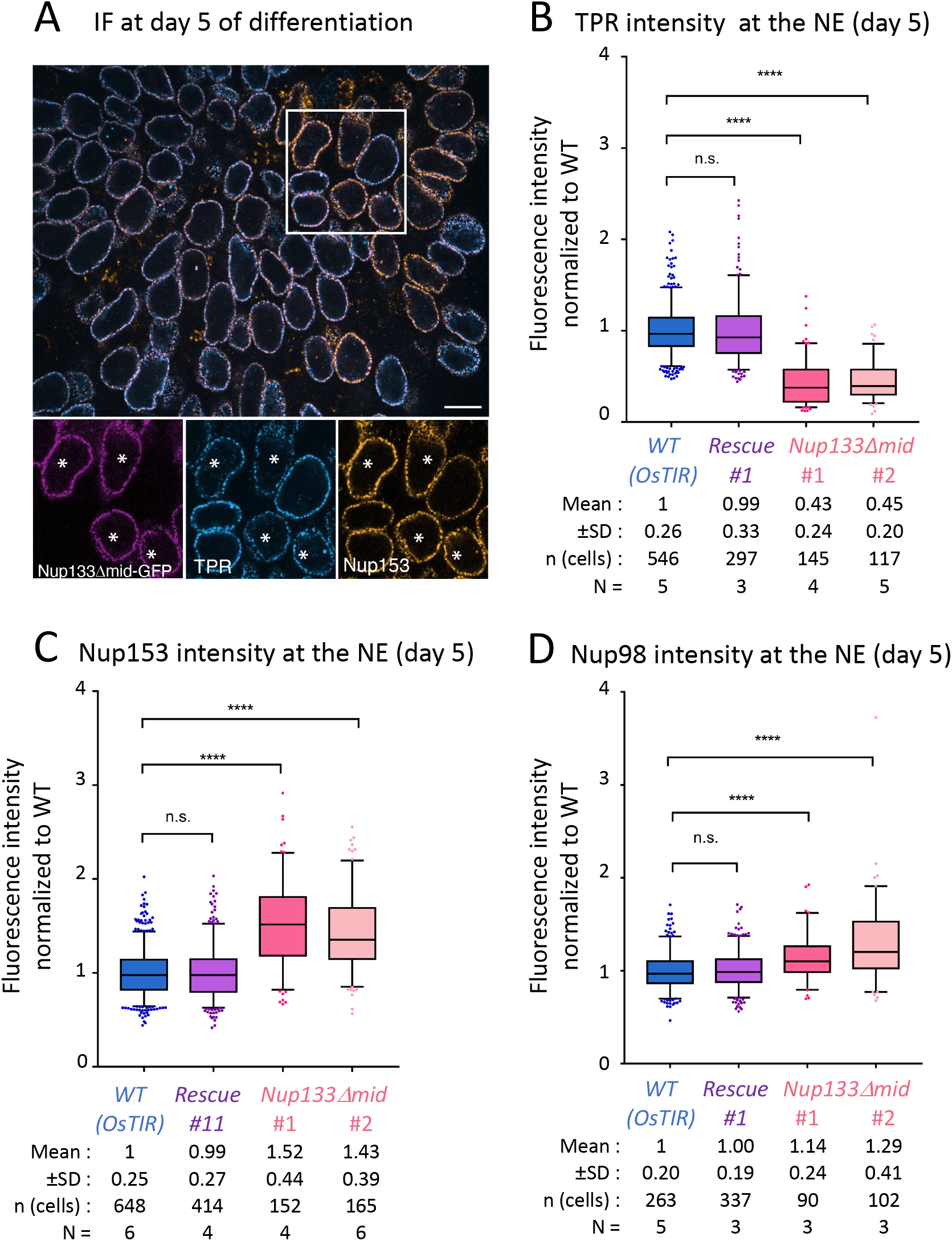
*Nup133∆mid* cells display a nuclear basket assembly defect at the neuronal progenitor stage. **A**. Representative image (single Z-section) of Tpr and Nup153 immunofluorescence of *WT (OsTIR)* mixed with *Nup133∆mid* cells (indicated by a *) at day 5 of differentiation. Scale bar, 10μm. **B, C, D**. Quantification of Tpr (B), Nup153 (C) and Nup98 (D) fluorescence intensity at the nuclear envelope, presented as box-plots. Values were normalized to the *WT (OsTIR)* in each field. Standard deviation (SD), number of analyzed cells (n) and of experiments (N) are indicated. ****: p-value<0.0001, n.s.: non-significant in Mann-Whitney test.

### Nup133-dependent gene regulation and nuclear basket assembly can be uncoupled

Having identified a critical function for the middle domain of Nup133 in gene regulation, we next aimed to determine the levels of Nup133 required for this process. We therefore established *Nup133-degron* cell lines that allow auxin-mediated degradation of a GFP-tagged allele of *Nup133* in an OsTIR-expressing mESC line (Gonzalez-Estevez, Verrico et al., (2021), see Materials and Methods and ***Figure S2B***).

The resulting *Nup133-degron* cell lines maintained normal *Nup133* mRNA expression during differentiation (***Figure S3A***), but actual Nup133 protein levels (without Auxin treatment) were only ∼25% of that found in *WT* cells (***Figure 4A and S3B***). This could be due to leaky OsTIR-induced degradation as previously reported (Mendoza-Ochoa et al., 2019; Yesbolatova et al., 2020), decreased stability of the tagged nucleoporin, or impaired export or translation of its mRNA. Nevertheless, these cells properly differentiated in the absence of auxin (***Figures 4C, S3D*** and ***S3E***). As anticipated, addition of auxin to the medium throughout differentiation led to a *Nup133*^*-/-*^*-*like phenotype: normal growth at the pluripotent state but massive cell death in neuronal differentiation (***Figures 4C, S3C*** and ***S3D***).

**Figure 4.**
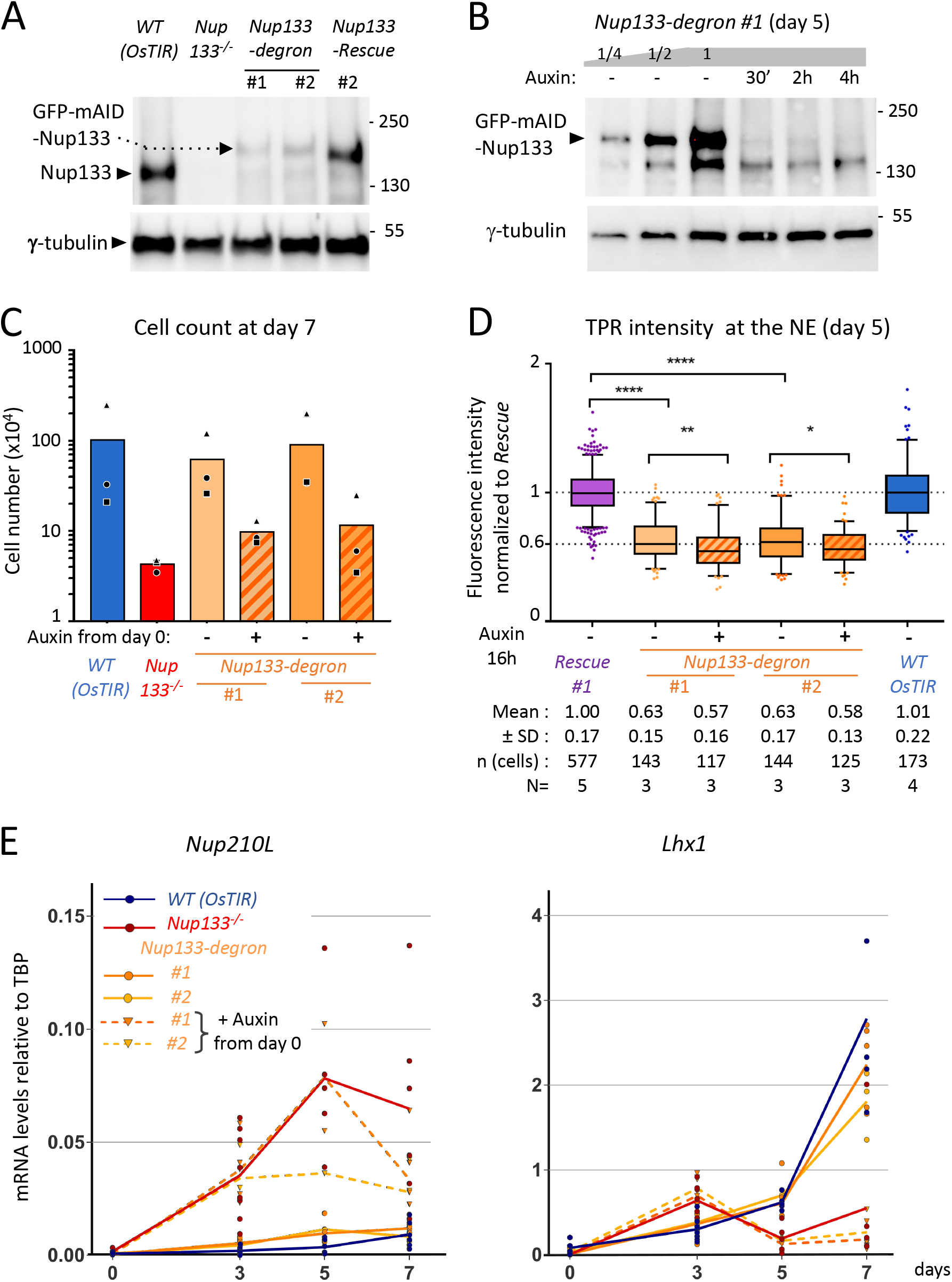
Nuclear basket integrity and gene regulation are uncoupled in *Nup133-degron* cells. **A, B**. Western blot showing the levels of endogenous Nup133 or its GFP-mAID-(*degron*) or GFP-(*Rescue*) tagged forms in the indicated cell lines at day 5 of differentiation. In **B**, *Nup133-degron* cells at day 5 of differentiation were either treated with ethanol (used as solvent for auxin; -) or with auxin for the indicated time. 1/2 and 1/4 dilutions of the non-treated *Nup133-degron* extract were also loaded. γ-tubulin is used as loading control. Molecular weights are indicated (kilodaltons). **C**. Cell counts at day 7 of neuroectodermal differentiation (n=3). Cells were seeded at 0.85 × 10^4^ cells/cm^2^. The graph represents the average of 3 independent experiments, each represented by a distinct label. **D**. Quantification of Tpr fluorescence intensity at the nuclear envelope in *Nup133-degron* cells treated (+) or not (-) for 16 h with auxin at day 5 of differentiation, presented as box-plots. Values were normalized to the *Nup133-Rescue* in each field. Standard deviation (SD), number of analyzed cells (n) and of experiments (N) are indicated. ****: p-value<0.0001; **: p-value<0.01; *: p-value<0.05; n.s.: non-significant in Mann-Whitney test. **E**. mRNA levels of *Lhx1* and *Nup210L* were quantified by RT-qPCR in *Nup133-degron* cells treated (dotted lines) or not (continuous lines) with auxin from day 0 on. The graph corresponds to 2-7 independent experiments for each cell line (represented by distinct labels).

Importantly, the lower Nup133 levels observed in the degron cell lines in the absence of auxin was not accompanied by the altered expression of *Nup210L* or *Lhx1* during neuronal differentiation (***Figure 4E***). In contrast, more extensive depletion of Nup133 upon continuous auxin treatment mimicked the effect of *Nup133* inactivation on these genes (***Figure 4E***).

Quantification of the nuclear basket protein Tpr in the *Nup133-degron* cell lines revealed that, even in the absence of auxin, Tpr levels at the nuclear envelope were already reduced to ∼60% of the WT levels both in pluripotent cells and in neuronal progenitors (***Figures 4D*** and ***S4A***). In contrast, the levels of Nup98 were not reduced in *Nup133-degron* cells (***Figure S4C***), consistent with a largely unaltered NPC density and a specific alteration of the nuclear basket. The minor - i.e., less than 10% - increase in Nup98 intensity observed in one of the two cell lines (*Nup133-degron* #1) may reflect modest clonal-dependent variations of NPC density. Finally, the levels of Nup153 were very mildly increased only in the *Nup133-degron* #1 cells, with a similar trend in both undifferentiated and differentiated cells (***Figures S4B*** and ***S4D***). These results indicate that a 4-fold reduction of Nup133 protein levels in untreated *Nup133-degron cells* is sufficient to severely impair Tpr recruitment or stabilization at nuclear pores, without major additional impact on NPC density.

Finally, although most of the GFP-mAID-Nup133 protein was already degraded after 30 minutes of auxin treatment in differentiated cells (***Figure 4B***), a 16 to 24h auxin treatment of these cells only led to a modest additional decrease of Tpr levels at the nuclear envelope compared to the untreated *Nup133-degron* cells (***Figures 4D*** and ***S4A***).

Overall, these results thus demonstrated that a correct Nup133 stoichiometry is critical for nuclear basket assembly, yet is not required for cell viability or gene regulation upon neuroectodermal differentiation. Taken together these data also reveal that a properly assembled nuclear basket at all NPCs is not required to regulate the expression of Nup133-target genes.

### *Nup210L* mRNA levels rapidly increase in response to Nup133 or Seh1 depletion

We next aimed to determine if the altered expression of *Nup210L* and *Lhx1* was specific for Nup133 or was shared by other Y-complex constituents. In view of the requirement for Seh1 in global NPC assembly, distinct from the specific basket assembly defect of *Nup133* mutant cells (Gonzalez-Estevez, Verrico et al., 2021), we chose to assess its implication at early stages of differentiation. Despite a very extensive death of the *Seh1*^*-/-*^ mutant cells upon differentiation, (Gonzalez-Estevez, Verrico et al., 2021), we could recover some mRNAs from these cells at day 3 of differentiation. As observed for *Nup133*^*-/-*^ mESCs (***Figure 1A***), *Seh1*^*-/-*^ cells properly repressed pluripotency markers and were able to induce early differentiation markers (***Figure S5A)***. Importantly, mRNA levels of both *Nup210L* and *Lhx1* were aberrantly increased in *Seh1*^*-/-*^ cells at day 3 of differentiation as also observed in differentiating *Nup133*^*-/-*^ cells (***Figure 5A***).

**Figure 5.**
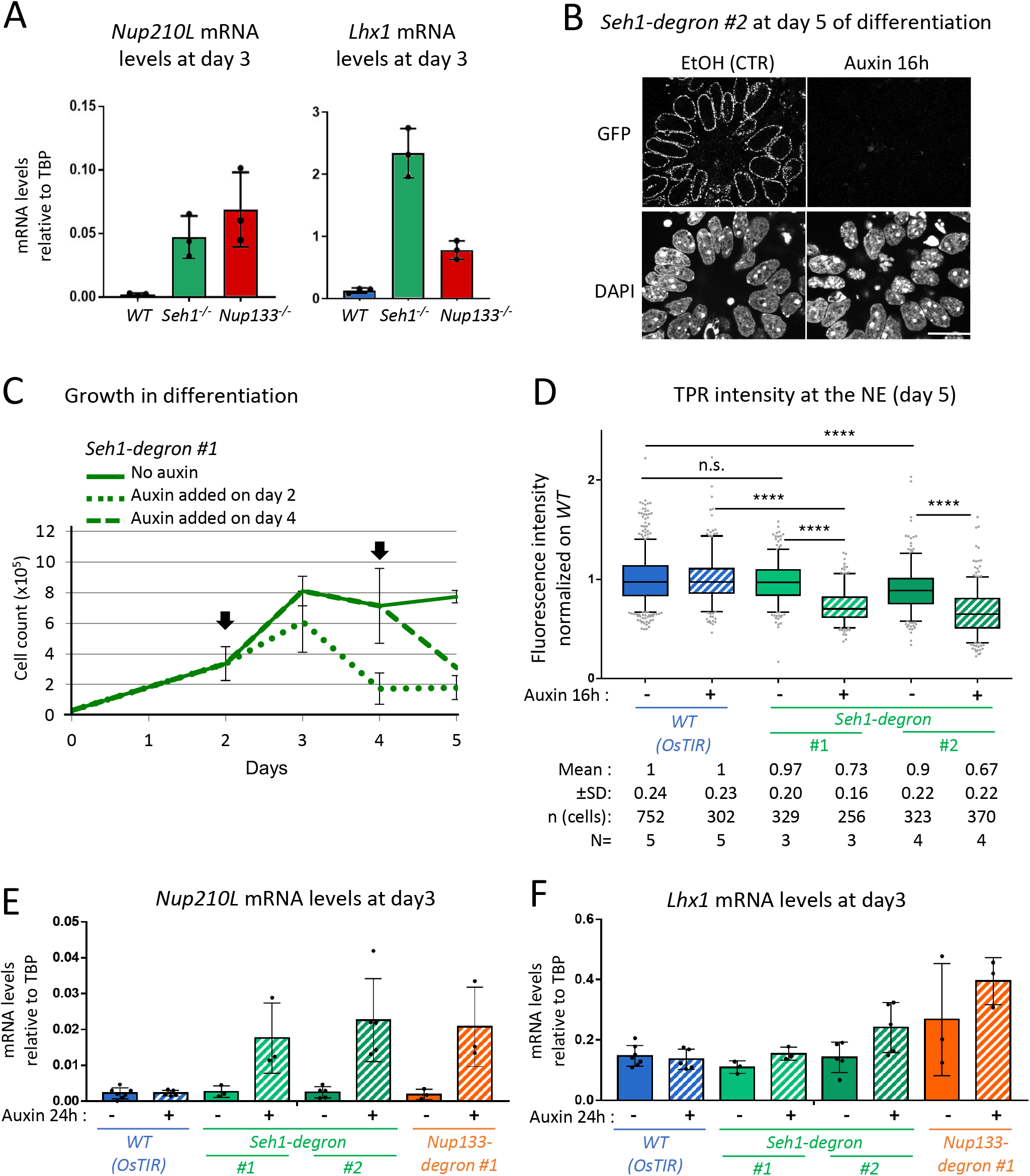
Seh1 depletion leads to *Lhx1* and *Nup210L* misregulation and altered NPC density in neuronal progenitors. **A**. Expression levels of *Lhx1* and *Nup210L* was assessed by RT-qPCR at day 3 of neuroectodermal differentiation in *WT* (HM1), *Seh1*^*-/-*^ (#1) and *Nup133*^*-/-*^ (#14) cells (n=3). The graph represents the average and standard deviation of 3 independent experiments, each represented by a dot. **B**. Immunofluorescence analysis of Seh1-mAID-GFP expressing cells (*Seh1-degron* #2) at day 5 of differentiation. Cells treated with EtOH or auxin for the last 16h were fixed and stained with DAPI. Scale bar, 20µm. **C**. Growth curve obtained from cell counts (n=3) in neuroectodermal differentiation of *Seh1-degron* cells treated when indicated with auxin from day 2 or day 4 on. Cells were seeded at 0.85 × 10^4^ cells/cm^2^. Mean and standard deviations are shown. **D**. Quantification of Tpr intensity at the NE at day 5 of differentiation, in the indicated cell lines. Cells were treated (+) or not (-) with auxin during the previous 16h. Values are normalized to *WT (OsTIR)* and presented as box-plots. Standard deviation (SD), number of analyzed cells (n) and of experiments (N) are indicated. ****: p-value<0.0001; n.s.: non-significant in Mann-Whitney test. **E, F**. Expression of *Lhx1* and *Nup210L* was assessed by RT-qPCR at day 3 of differentiation in cells treated with auxin (+) or EtOH (-) for the last 24 h.

The low viability of *Seh1*^*-/-*^ cells upon differentiation (Gonzalez-Estevez, Verrico et al., 2021) did not allow us to perform quantitative immunofluorescence studies at the differentiated stage. We therefore established and validated new *Seh1-degron* cell lines (see Materials and Methods ***and Figure S2C***), in which the C-terminally-tagged form of Seh1 was properly and homogeneously expressed upon differentiation (***Figures 5B, S5B*** and ***S5C***). In the resulting *Seh1-degron* cells, 24h addition of auxin at day 2 or 4 of differentiation led to impaired viability of the cells (***Figure 5C***). This indicates that Seh1 is not solely required at the early onset of neuronal progenitor differentiation, but also for the proper growth or viability of the progenitors themselves. Analyses of nuclear pore assembly in *Seh1-degron*-derived neuronal progenitors (at day 5 of differentiation) did not reveal major defects in the absence of auxin (***Figure 5D***). In contrast, a 16h-treatment with auxin led to a ∼30% decrease of both Tpr and Nup98 intensities at the nuclear envelope of neuronal progenitors compared to the control cells (***Figures 5D*** and ***S4E***). This suggests a global decrease in pore number upon Seh1 depletion, comparable to the observations previously made in pluripotent *Seh1*^*-/-*^ and *GFP-mAID-Seh1* mESCs (Gonzalez-Estevez, Verrico et al., 2021). Note that unlike Tpr and Nup98, Nup153 levels were not altered in auxin-treated *Seh1-degron* cells, suggesting, as also observed in *Nup133∆mid* cells, an increased stoichiometry of Nup153 per NPC (***Figure S4F***). We also note that a 24h auxin treatment, applied to *Seh1-mAID-GFP* cells at day 2 of neuronal differentiation was sufficient to cause an important increase in *Nup210L* mRNA levels (***Figure 5E***). Likewise, a 24h auxin treatment of *Nup133-degron* cells induced *Nup210L* expression (***Figure 5E***). In contrast, 24h of auxin treatment did not lead to an altered regulation of Lhx1 in *Seh1-degron* or *Nup133-degron* cells at day 3 (***Figure 5F***). Together, these data indicate that *Nup210L* and *Lhx1* are shared downstream targets of Nup133 and Seh1, with *Nup210L* appearing to be a gene induced early upon loss of these Y-complex Nups. On the other hand, *Lhx1* seems to need a prolonged depletion of these Y-complex Nups to become misregulated, indicating that it is likely an indirect target of Y-complex Nup-dependent regulations.

## CONCLUSION

In this study, we have shown in differentiating mESCs that a subset of genes is deregulated in the absence of Nup133. Although neuronal progenitors lacking either Nup133 or just its middle domain share a defect in nuclear basket assembly and altered regulation of *Nup210L* and *Lhx1*, these two phenotypes can be uncoupled. Indeed, these two genes were not misregulated in our *Nup133-degron* cell lines that display a constitutive nuclear basket assembly defect, and conversely, they were both similarly misregulated in *Seh1-deficient* cells in which nuclear basket assembly is not specifically altered. As the untreated *Nup133-degron* cell lines exhibit lower Nup133 protein levels than the control cell lines, our data further argue that a limited amount of Nup133 is sufficient to keep *Nup210L* repressed in differentiating mESCs and to induce the proper and timely expression of *Lhx1*. Because this function in gene regulation is shared by Nup133 and Seh1, two physically distant members of the Y-complex, it likely involves the whole Y-complex rather than each of these two individual subunits.

The rather short lag time (below 24h) between auxin-induced degradation of Nup133 or Seh1 and *Nup210L* activation suggests that there could be a direct contact between the Y-complex and the *Nup210L* genomic locus. Although the Y-complex is mainly visualized at NPCs to which it is stably anchored (Rabut et al., 2004), a diffuse fraction is also likely present in the nucleus, as previously described in HeLa cells (Morchoisne-Bolhy et al., 2015). Hence, Y-complex dependent gene regulation may take place either at NPCs or “off-pore”. Browsing available data of LaminB1-Dam-ID tracks (Peric-Hupkes et al., 2010), we noticed that the *Nup210L* locus is adjacent to a lamin-associated domain (LAD) in mESCs and neural progenitor cells. A location near the nuclear periphery would be consistent with a regulation of *Nup210L* taking place at NPCs. In line with this hypothesis, Seh1 was shown to recruit to the NPC the transcription factor Olig2 and the chromatin remodeler Brd7 to promote the expression of differentiation genes in oligodendrocytes (Liu et al., 2019). Additionally, Nup133 was proposed to promote the expression of *Myc* in cancer cells by anchoring its superenhancer to the NPCs (Scholz et al., 2019).

This Y-complex-mediated gene regulation may also involve epigenetic mechanisms, as reported for Nup153, which interacts with PRC1 to repress developmental genes (Jacinto et al., 2015). Along these lines, it is noteworthy that human *NUP210L*, initially thought to be a testis-specific gene, was found to be expressed in prefrontal cortex neurons of some individuals. This regulation was linked to the epigenetic, allele-specific activation of *NUP210L*, namely the deposition of the permissive histone mark H3K4me3 at its promoter (Gusev et al., 2019). In addition, another epigenetic mechanism, the DNA methylation state of *Nup210L*, has been linked to psychologic development disorders in patients carrying a hemizygous 22q11.2 microdeletion (Starnawska et al., 2017). Considering its possible link with normal or pathological cognitive abilities, the mechanisms and consequences of Nup133- and Seh1-dependent *Nup210L* activation warrant further investigations.

## MATERIALS AND METHODS

### mESCs culture and neuroectodermal differentiation

Cell lines used in this study are listed in ***Table S1*** and were grown as previously described (Gonzalez-Estevez, Verrico et al., 2021). Briefly, mESCs were grown at 37°C and 5% CO2 on Mitomycin-C inactivated feeder cells (DR4-mouse embryonic fibroblast) plated on 0.1% gelatin (Sigma-Aldrich) in serum/leukemia inhibitory factor (LIF, ESGRO, Millipore)-containing stem cell medium.

The neuroectodermal differentiation protocol used in this study was adapted from (Abranches et al., 2009; Ying et al., 2003), as previously described (Gonzalez-Estevez, Verrico et al., 2021). Briefly, following trypsinization and feeder removal, mESCs were resuspended in N2B27 medium [Neurobasal, DMEM-F12, 7.5% BSA, N2 supplement, B27 supplement, Pen/Strep, L-glutamin, ß-mercaptoethanol] and plated at a density of ∼ 0.85×10^4^ or 3×10^4^ cells/cm^2^ on gelatin-coated dishes (day 0). Medium was changed every day from day 2 on. To stimulate neuronal differentiation, 1μM RA (all-trans-Retinoic acid, Sigma) was added to the medium for 24h on day 2.

For annexin V/propidium iodide (PI) apoptosis/viability assays, cells were trypsinized, counted, and 10^5^ cells were centrifuged at 400 x *g* for 3 minutes. Cells were resuspended in 500µL of Annexin V binding buffer (ab14084, Abcam) and incubated with 1µL annexin V-Cy5 (ab14147, Abcam) and 10 µg/mL propidium iodide for 5min at room temperature in the dark. Cells were then analyzed by flow cytometry using a CyanADP Cytomation (Beckman-Coulter), using SS (side-scatter) and FS (forward scatter) to remove debris and exclude cell doublets, and 488 nm and 635 nm excitation lasers. At least 10.000 cells were acquired and data were then processed using the Summit software.

To induce degradation of the GFP-mAID-Nup133 (in *Nup133-degron* cells) and Seh1-mAID-GFP (in *Seh1-degron* cells), 500µM auxin (Sigma-Aldrich) was added to the medium (from a 280 mM stock in EtOH). The same final concentration of EtOH was added for control experiments.

### RNA-sequencing

RNAs were extracted from 3 independent *Nup133*^*-/-*^ clones (KO#1, HM1-derived *Nup133*^*-/-*^ #14; KO#2, HM1-derived *Nup133*^*-/-*^ #19; KO#3, blastocyst-derived #319 *Nup133*^*merm/merm*^ mESCs), and from 3 isogenic control samples (WT#1; HM1; WT#2, HM1; WT#3, blastocyst-derived #1AA *Nup133*^*+/+*^) (See ***Table S1***). Library preparation and Illumina sequencing were performed at the Ecole normale superieure genomics core facility (Paris, France). Messenger (polyA+) RNAs were purified from 1 µg of total RNA using oligo(dT). Libraries were prepared using the strand specific RNA-Seq library preparation TruSeq Stranded mRNA kit (Illumina). Libraries were multiplexed by 9 on 2 flowcells. Two 75 bp single read sequencing runs were performed on a NextSeq 500 device (Illumina). A mean of 53.14 ± 14.72 million passing Illumina quality filter reads was obtained for each of the 18 samples.

The analyses were performed using the Eoulsan pipeline version 2.0-alpha7 (Jourdren et al., 2012), including read filtering, mapping, alignment filtering, read quantification, normalization and differential analysis: Before mapping, poly N read tails were trimmed, reads ≤40 bases were removed, and reads with quality mean ≤30 were discarded. Reads were then aligned against the *Mus musculus* genome from Ensembl version 81 using STAR (version 2.4.0k)(Dobin et al., 2013). Alignments from reads matching more than once on the reference genome were removed using Java version of samtools (Li et al., 2009). To compute gene expression, *Mus musculus* GFF genome annotation version 81 from Ensembl database was used. All overlapping regions between alignments and referenced exons were counted and aggregated by genes using the HTSeq-count algorithm (Anders et al., 2015). A first analysis revealed that one of the samples (KO#2 day2) featured an abnormally high level of ribosomal transcripts; this dataset was therefore excluded from subsequent analyses.

The rest of the analysis was carried out using the bioinformatics software R (R v4.1.2 (2021.11.01)), and open access packages, using the publicly available bioinformatics course DIYtranscriptomics.com (Berry et al., 2021). Mapped raw counts were transformed in counts per million (cpm) using the cpm function from the EdgeR package (v3.34.1). We filtered the genes that had a log2(cpm) above 1 for more than 3 samples, and then normalized their cpm using the TMM method (Robinson and Oshlack, 2010). The mean-variance relationship of the filtered normalized data was modeled by voom transformation, and a linear model was fitted to the data using the lmfit function from the limma package (v3.48.3). Bayesian statistics for the chosen pair-wise comparisons (average KO expression compared to average WT expression for each time point) were then calculated using the eBayes function from limma, and adjusted with the BH correction. An exhaustive list of differentially expressed genes (p. value<0.05 and |logFC|>1.5) was pulled-out using the decideTests function. Plots in ***Figures 1D and S1A*** were generated in R using ggplot2 (v.3.3.5).

The RNASeq gene expression data and raw fastq files are available on the GEO repository (https://www.ncbi.nlm.nih.gov/geo/query/acc.cgi?acc=GSE218080) under accession number: GSE218080.

### Transfection and CRISPR/Cas9 genome editing

mESCs were transfected as previously described (Souquet et al., 2018) using Lipofectamin 2000 (Invitrogen) according to manufacturer instructions. To establish the *Nup133-Rescue, Nup133-∆mid, Nup133-degron* and *Seh1-degron* cell lines, 5·10^5^ mESCs were co-transfected with 3µg of a plasmid directing the expression of a gRNA and high fidelity (HF) Cas9 fused to mCherry and with 3 µg of DNA sequences of interest flanked by homology directed repair arms (PCR product or linearized plasmid, see ***Figure S2***). Plasmids are listed in ***Table S2***, gRNAs designed using the Benchling website (https://benchling.com) are listed in ***Table S3***, and PCR primers used to generate homology-directed repair templates are listed in ***Table S4***. 3 days after transfection, GFP-positive cells were FACS-sorted to select for cells expressing the tagged nucleoporin and plated on culture dishes. Individual clones were picked 6-7 days after sorting and characterized using immunofluorescence, western blot, PCR on genomic DNA and sequencing. Ploidy was assessed using chromosome spreads. DAPI staining was used to ensure lack of major contamination by mycoplasma. Cell line characteristics are summarized in ***Table S1***.

### RT-qPCR

RNA extraction was performed using NucleoSpin RNAII isolation kit (Mascherey-Nagel) according to the manufacturer’s instructions. Reverse-transcription (RT) was done with the transcriptase inverse Superscript II (Invitrogen) and random hexamers (Amersham Pharmacia), using at least 150 ng of RNA per sample. Real-time quantitative PCR was performed with SybrGreen reagents (Applied Biosystems) on a LightCycler 480 instrument (Roche Life Sciences). All mRNA level results are presented as relative to the TATA-binding protein (TBP) mRNA levels. qPCR primers used in this study are listed in supplementary ***Table S5***.

### Western blots analyses

Whole cell lysate preparations and western blot analyses were performed as previously described (Gonzalez-Estevez, Verrico et al., 2021), using 4-10% SDS-PAGE gels (Mini-Protean TGX Stain free precast gels, Bio-rad) and nitrocellulose membranes (GE healthcare). Incubations with primary antibodies were carried overnight at 4°C. Signals from HRP-conjugated secondary antibodies were detected by enhanced chemiluminescence (SuperSignal® Pico or Femto, ThermoScientific) using ChemiDoc (Biorad). Antibodies used in this study are listed in ***Table S6***.

### Immunofluorescence and quantification of nucleoporin intensity at the nuclear envelope

Cells grown on glass coverslips coated with 0.1% gelatin were fixed for 20 minutes in 3% paraformaldehyde (VWR, Radnor, PA) (resuspended in PBS and brought to pH 8.0 with NaOH), permeabilized 30 minutes in H-Buffer (PBS, BSA 1%, Triton 0.2%, SDS 0.02%) and incubated with the primary and secondary antibodies for 1h at room temperature in H-Buffer, with washes in H-Buffer in-between. Antibodies used in this study are listed in supplementary ***Table S5***. Coverslips were then incubated 5 min with DAPI (Sigma, 280nM in PBS) and mounted using Vectashield (Vector, Maravai Life Sciences, San Diego, CA). Images were acquired on a DMI8 microscope (Leica), equipped with a CSU-W1 spinning-diskhead (Yokogawa, Japan) and 2 Orca-Flash 4 V2+ sCMOS cameras (Hamamatsu), using 100x/1.4 oil objectives Quantification of nucleoporin intensities at the nuclear envelope (NE) was performed essentially as described (Souquet et al., 2018), by mixing the cell line of interest with a reference cell line, either the *WT (OsTIR)* cell line or the *Nup133-Rescue* line, as indicated. For each field, we measured the mean intensity of 8-pixel-thick lines drawn on the nuclear rims and of a background area. After subtraction of the background, the NE intensity value obtained for each cell was normalized to the mean value obtained for the reference cells acquired in the same field. Box plots were generated using GraphPad Software: each box encloses 50% of the normalized values obtained, centered on the median value. The bars extend from the 5^th^ to 95^th^ percentiles. Values falling outside of this range are displayed as individual points. Statistical analyses were performed using unpaired nonparametric Mann-Whitney tests. p<0.0001=****, p<0.001=***, p<0.01=**, p<0.05=*.

## Acknowledgements

We are grateful to Vedrana Andric, Alessandro Berto, Marta Boira and Salomé Neuvendel for help in mESCs culture, differentiation, cell line establishment, or western blot analyses. We thank Charlène Boumendil and Pierre Therizols for helpful discussions, and Benoit Palancade, Roger Karess, Charlène Boumendil and the Doye lab members for critical reading of the manuscript. We also acknowledge the ImagoSeine core facility of the Institut Jacques Monod, for help with cell sorting, FACS analyses, and spinning disk imaging.

## AUTHOR CONTRIBUTIONS

C.O., A.V., B.S., and V.D. conceived and designed the experiments.

C.O., A.V., S.P., B.S., F.C. and S.B. performed the experiments.

C.O., A.V., S.P., L.J. and V.D. analyzed the data

C.O., A.V., and V.D. wrote the manuscript with contribution of L. J. for the method section.

## REFERENCES

Abranches, E., Silva, M., Pradier, L., Schulz, H., Hummel, O., Henrique, D., and Bekman, E. (2009). Neural differentiation of embryonic stem cells in vitro: A road map to neurogenesis in the embryo. PLoS One 4, e6286.

Aksenova, V., Smith, A., Lee, H., Bhat, P., Esnault, C., Chen, S., Iben, J., Kaufhold, R., Yau, K.C., Echeverria, C., et al. (2020). Nucleoporin TPR is an integral component of the TREX-2 mRNA export pathway. Nat. Commun. 11, 1–13.

Anders, S., Pyl, P.T., and Huber, W. (2015). HTSeq-A Python framework to work with high-throughput sequencing data. Bioinformatics 31, 166–169.

Berry, A.S.F., Amorim, C.F., Berry, C.L., Syrett, C.M., English, E.D., and Beiting, D.P. (2021). An open-source toolkit to expand bioinformatics training in infectious diseases. MBio 12, 1–6.

Berto, A., Yu, J., Morchoisne-Bolhy, S., Bertipaglia, C., Vallee, R., Dumont, J., Ochsenbein, F., Guerois, R., and Doye, V. (2018). Disentangling the molecular determinants for Cenp-F localization to nuclear pores and kinetochores. EMBO Rep. 19, e44742.

Boumendil, C., Hari, P., Olsen, K.C.F., Acosta, J.C., and Bickmore, W.A. (2019). Nuclear pore density controls heterochromatin reorganization during senescence. Genes Dev. 33, 144–149.

Braun, D.A., Lovric, S., Schapiro, D., Schneider, R., Marquez, J., Asif, M., Hussain, M.S., Daga, A., Widmeier, E., Rao, J., et al. (2018). Mutations in multiple components of the nuclear pore complex cause nephrotic syndrome. J. Clin. Invest. 128, 4313–4328.

Buchwalter, A., Kaneshiro, J.M., and Hetzer, M.W. (2019). Coaching from the sidelines: the nuclear periphery in genome regulation. Nat. Rev. Genet. 20, 39–50.

Buchwalter, A.L., Liang, Y., and Hetzer, M.W. (2014). Nup50 is required for cell differentiation and exhibits transcription-dependent dynamics. Mol. Biol. Cell 25, 2472–2484.

Costello, I., Nowotschin, S., Sun, X., Mould, A.W., Hadjantonakis, A.K., Bikoff, E.K., and Robertson, E.J. (2015). Lhx1 functions together with Otx2, Foxa2, and Ldb1 to govern anterior mesendoderm, node, and midline development. Genes Dev. 29, 2108–2122.

D’Angelo, M.A., Gomez-Cavazos, J.S., Mei, A., Lackner, D.H., and Hetzer, M.W. (2012). A Change in Nuclear Pore Complex Composition Regulates Cell Differentiation. Dev. Cell 22, 446–458.

Delay, B.D., Corkins, M.E., Hanania, H.L., Salanga, M., Deng, J.M., Sudou, N., Taira, M., Horb, M.E., and Miller, R.K. (2018). Tissue-specific gene inactivation in xenopus laevis: Knockout of lhx1 in the kidney with CRISPR/Cas9. Genetics 208, 673–686.

Dobin, A., Davis, C.A., Schlesinger, F., Drenkow, J., Zaleski, C., Jha, S., Batut, P., Chaisson, M., and Gingeras, T.R. (2013). STAR: Ultrafast universal RNA-seq aligner. Bioinformatics 29, 15–21.

Doucet, C.M., and Hetzer, M.W. (2010). Nuclear pore biogenesis into an intact nuclear envelope. Chromosoma 119, 469–477.

Dultz, E., Wojtynek, M., Medalia, O., and Onischenko, E. (2022). The Nuclear Pore Complex: Birth, Life, and Death of a Cellular Behemoth. Cells 11, 1–28.

Festuccia, N., Owens, N., Papadopoulou, T., Gonzalez, I., Tachtsidi, A., Vandoermel-Pournin, S., Gallego, E., Gutierrez, N., Dubois, A., Cohen-Tannoudji, M., et al. (2019). Transcription factor activity and nucleosome organization in mitosis. Genome Res. 29, 250–260.

Fujita, A., Tsukaguchi, H., and Koshimizu, E. (2018). Homozygous Splicing Mutation in NUP133 Causes Galloway–Mowat Syndrome Atsushi. Ann. Neurol. 84, 814–828.

Gonzalez-Estevez, A., Verrico, A., Orniacki, C., Reina-San-Martin, B., and Doye, V. (2021). Integrity of the short arm of the nuclear pore Y-complex is required for mouse embryonic stem cell growth and differentiation. J. Cell Sci. 134, jcs258340.

Gusev, F.E., Reshetov, D.A., Mitchell, A.C., Andreeva, T. V., Dincer, A., Grigorenko, A.P., Fedonin, G., Halene, T., Aliseychik, M., Goltsov, A.Y., et al. (2019). Epigenetic-genetic chromatin footprinting identifies novel and subject-specific genes active in prefrontal cortex neurons. FASEB J. 33, 8161–8173.

Harel, A., Orjalo, A. V., Vincent, T., Lachish-Zalait, A., Vasu, S., Shah, S., Zimmerman, E., Elbaum, M., and Forbes, D.J. (2003). Removal of a single pore subcomplex results in vertebrate nuclei devoid of nuclear pores. Mol. Cell 11, 853–864.

Hezwani, M., and Fahrenkrog, B. (2017). The functional versatility of the nuclear pore complex proteins. Semin. Cell Dev. Biol. 68, 2–9.

Huang, Y., Chavez, L., Chang, X., Wang, X., Pastor, W.A., Kang, J., Zepeda-Martínez, J.A., Pape, U.J., Jacobsen, S.E., Peters, B., and Rao, A. (2014). Distinct roles of the methylcytosine oxidases Tet1 and Tet2 in mouse embryonic stem cells. Proc. Natl. Acad. Sci. U.S.A. 111:1361–1366.

Jacinto, F. V., Benner, C., and Hetzer, M.W. (2015). The nucleoporin Nup153 regulates embryonic stem cell pluripotency through gene silencing. Genes Dev. 29, 1224–1238.

Jourdren, L., Bernard, M., Dillies, M.A., and Le Crom, S. (2012). Eoulsan: A cloud computing-based framework facilitating high throughput sequencing analyses. Bioinformatics 28, 1542– 1543.

Jühlen, R., and Fahrenkrog, B. (2018). Moonlighting nuclear pore proteins: tissue-specific nucleoporin function in health and disease. Histochem. Cell Biol. 150, 593–605.

Krull, S., Dörries, J., Boysen, B., Reidenbach, S., Magnius, L., Norder, H., Thyberg, J., and Cordes, V.C. (2010). Protein Tpr is required for establishing nuclear pore-associated zones of heterochromatin exclusion. EMBO J. 29, 1659–1673.

Li, H., Handsaker, B., Wysoker, A., Fennell, T., Ruan, J., Homer, N., Marth, G., Abecasis, G., and Durbin, R. (2009). The Sequence Alignment/Map format and SAMtools. Bioinformatics 25, 2078– 2079.

Liu, Z., Yan, M., Liang, Y., Liu, M., Zhang, K., Shao, D., Jiang, R., Li, L., Wang, C., Nussenzveig, D.R., et al. (2019). Nucleoporin Seh1 Interacts with Olig2/Brd7 to Promote Oligodendrocyte Differentiation and Myelination. Neuron 102, 587–601.

Lupu, F., Alves, A., Anderson, K., Doye, V., and Lacy, E. (2008). Nuclear Pore Composition Regulates Neural Stem/Progenitor Cell Differentiation in the Mouse Embryo -Supplemental-. Dev. Cell 14, 831–842.

McCloskey, A., Ibarra, A., and Hetzer, M.W. (2018). Tpr regulates the total number of nuclear pore complexes per cell nucleus. Genes Dev. 32, 1321–1331.

McMahon, R., Sibbritt, T., Salehin, N., Osteil, P., and Tam, P.P.L. (2019). Mechanistic insights from the LHX1-driven molecular network in building the embryonic head. Dev. Growth Differ. 61, 327–336.

Mendoza-Ochoa, G.I., Barrass, J.D., Terlouw, B.R., Maudlin, I.E., de Lucas, S., Sani, E., Aslanzadeh, V., Reid, J.A.E., and Beggs, J.D. (2019). A fast and tuneable auxin-inducible degron for depletion of target proteins in budding yeast. Yeast 36, 75–81.

Morchoisne-Bolhy, S., Geoffroy, M.C., Bouhlel, I.B., Alves, A., Audugé, N., Baudin, X., Van Bortle, K., Powers, M.A., and Doye, V. (2015). Intranuclear dynamics of the Nup107-160 complex. Mol. Biol. Cell 26, 2343–2356.

Ori, A., Banterle, N., Iskar, M., Andrés-Pons, A., Escher, C., Khanh Bui, H., Sparks, L., Solis-Mezarino, V., Rinner, O., Bork, P., et al. (2013). Cell type-specific nuclear pores: A case in point for context-dependent stoichiometry of molecular machines. Mol. Syst. Biol. 9, 648.

Peric-Hupkes, D., Meuleman, W., Pagie, L., Bruggeman, S.W.M., Solovei, I., Brugman, W., Gräf, S., Flicek, P., Kerkhoven, R.M., van Lohuizen, M., et al. (2010). Molecular Maps of the Reorganization of Genome-Nuclear Lamina Interactions during Differentiation. Mol. Cell 38, 603–613.

Rabut, G., Doye, V., and Ellenberg, J. (2004). Mapping the dynamic organization of the nuclear pore complex inside single living cells. Nat. Cell Biol. 6, 1114–1121.

Robinson, M.D., and Oshlack, A. (2010). A scaling normalization method for differential expression analysis of RNA-seq data. Genome Biol. 11, 1–9.

Scholz, B.A., Sumida, N., de Lima, C.D.M., Chachoua, I., Martino, M., Tzelepis, I., Nikoshkov, A., Zhao, H., Mehmood, R., Sifakis, E.G., et al. (2019). WNT signaling and AHCTF1 promote oncogenic MYC expression through super-enhancer-mediated gene gating. Nat. Genet. 51, 1723–1731.

Shawlot, W., Wakamiya, M., Kwan, K.M., Kania, A., Jessell, T.M., and Behringer, R.R. (1999). Lim1 is required in both primitive streak-derived tissues and visceral endoderm for head formation in the mouse. Development 126, 4925–4932.

Souquet, B., Freed, E., Berto, A., Andric, V., Audugé, N., Reina-San-Martin, B., Lacy, E., and Doye, V. (2018). Nup133 Is Required for Proper Nuclear Pore Basket Assembly and Dynamics in Embryonic Stem Cells. Cell Rep. 23, 2443–2454.

Starnawska, A., Hansen, C.S., Sparsø, T., Mazin, W., Olsen, L., Bertalan, M., Buil, A., Bybjerg-Grauholm, J., Bækvad-Hansen, M., Hougaard, D.M., et al. (2017). Differential DNA methylation at birth associated with mental disorder in individuals with 22q11.2 deletion syndrome. Transl. Psychiatry 7, e1221.

Subrini, J., and Turner, J. (2021). Y chromosome functions in mammalian spermatogenesis. Elife 10, 1–20.

Toda, T., Hsu, J.Y., Linker, S.B., Hu, L., Schafer, S.T., Mertens, J., Jacinto, F. V, Hetzer, M.W., and Gage, F.H. (2017). Nup153 Interacts with Sox2 to Enable Bimodal Gene Regulation and Maintenance of Neural Progenitor Cells. Cell Stem Cell 21, 618–634.

Vollmer, B., Lorenz, M., Moreno-Andrés, D., Bodenhöfer, M., De Magistris, P., Astrinidis, S.A., Schooley, A., Flötenmeyer, M., Leptihn, S., and Antonin, W. (2015). Nup153 Recruits the Nup107-160 Complex to the Inner Nuclear Membrane for Interphasic Nuclear Pore Complex Assembly. Dev. Cell 33, 717–728.

Walther, T.C., Alves, A., Pickersgill, H., Loïodice, I., Hetzer, M., Galy, V., Hülsmann, B.B., Köcher, T., Wilm, M., Allen, T., et al. (2003). The conserved Nup107-160 complex is critical for nuclear pore complex assembly. Cell 113, 195–206.

Wozniak, R.W., Bartnik, E., and Blobel, G. (1989). Primary structure analysis of an integral membrane glycoprotein of the nuclear pore. J. Cell Biol. 108, 2083–2092.

Yesbolatova, A., Saito, Y., Kitamoto, N., Makino-Itou, H., Ajima, R., Nakano, R., Nakaoka, H., Fukui, K., Gamo, K., Tominari, Y., et al. (2020). The auxin-inducible degron 2 technology provides sharp degradation control in yeast, mammalian cells, and mice. Nat. Commun. 11, 5701.

Ying, Q.L., Stavridis, M., Griffiths, D., Li, M., and Smith, A. (2003). Conversion of embryonic stem cells into neuroectodermal precursors in adherent monoculture. Nat. Biotechnol. 21, 183–186.

Zeng, H., Horie, K., Madisen, L., Pavlova, M.N., Gragerova, G., Rohde, A.D., Schimpf, B.A., Liang, Y., Ojala, E., Kramer, F., et al. (2008). An inducible and reversible mouse genetic rescue system. PLoS Genet. 4, e1000069.

