## Supplemental Figures S1-S5 and Supplemental Tables S1-S6 for "Y-complex nucleoporins independently contribute to nuclear pore assembly and gene regulation in neuronal progenitors"

### Inventory of Supplementary Material

**Supplemental Figures and Figure legend:**

**Figure S1, relative to Figure 1:** DEGs analysis in differentiating *Nup133*$^{-/-}$ cells.

**Figure S2, related to Materials and Methods and Tables S1, S3 and S4:** Schematics of CRISPR-Cas9-mediated cell line establishment via homologous recombination.

**Figure S3, relative to Figure 4:** Characterization of the *Nup133-degron* cell lines.

**Figure S4, related to Figures 3, B-D and Figure 4D:** Nucleoporin densities at the nuclear envelope in *Nup133-degron* and *Seh1-degron* cell lines.

**Figure S5, relative to Figure 5:** Characterization of the *Seh1*$^{-/-}$ and *Seh1-degron* cell lines.

**Supplemental Tables:**

**Table S1:** Cell lines used in this study

**Table S2:** Plasmids used in this study

**Table S3:** Sequences of gRNAs used in this study

**Table S4:** Primers used to generate homology-directed repair templates for Nup133- and Seh1-degrons

**Table S5:** qPCR primers used in this study

**Table S6:** Antibodies used in this study

**References cited in Supplemental Figures and Tables**

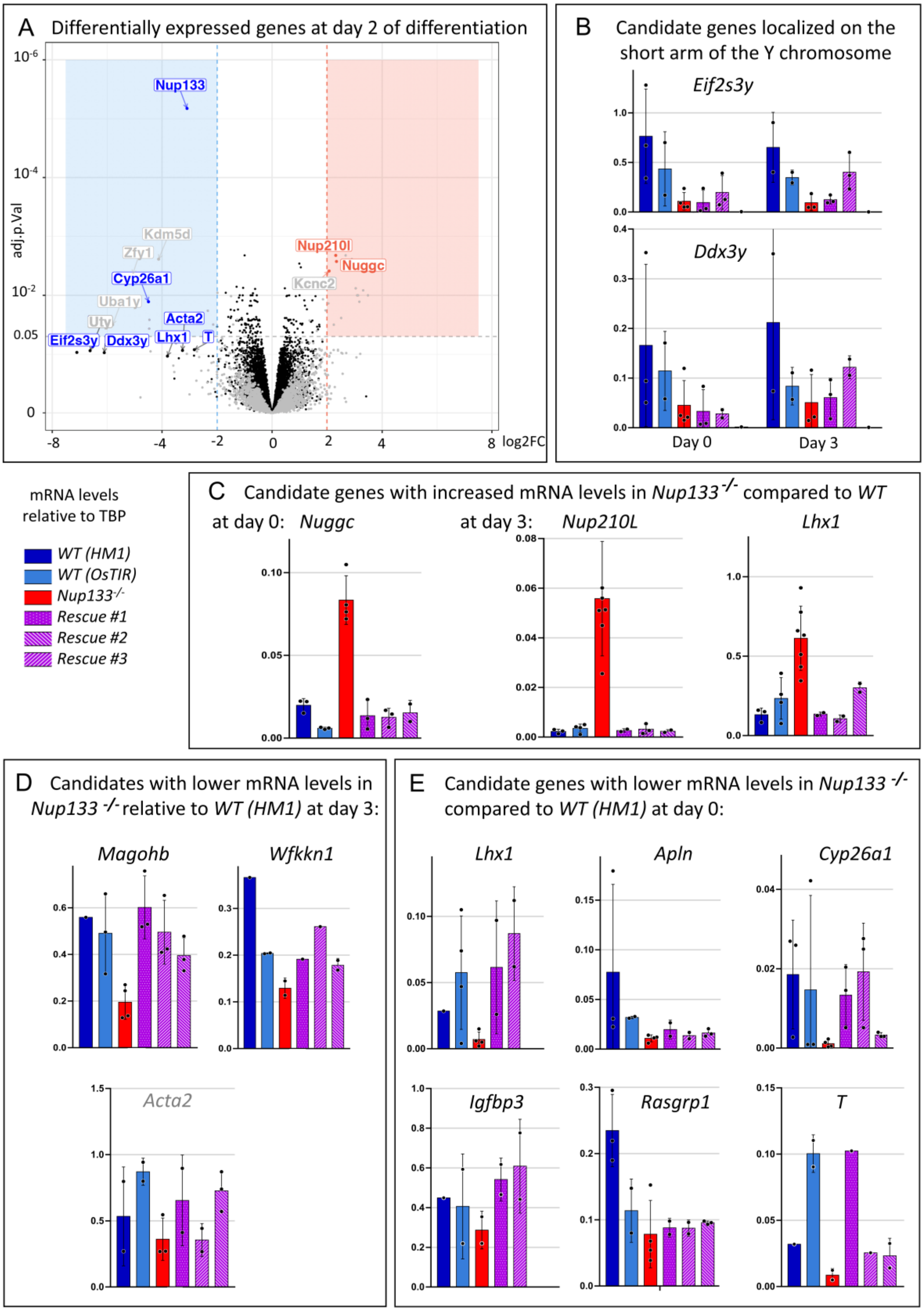

A Differentially expressed genes at day 2 of differentiation
adj.p.Val
log2FC
Nup133
Kdm5d
Zfy1
Cyp26a1
Uba1y
Uty
Acta2
Eif2s3y
Ddx3y
Lhx1
T
Nup210l
Nuggc
Kcnc2
B Candidate genes localized on the short arm of the Y chromosome
Eif2s3y
Ddx3y
Day 0
Day 3
mRNA levels relative to TBP
WT (HM1)
WT (OsTIR)
Nup133-/-
Rescue #1
Rescue #2
Rescue #3
C Candidate genes with increased mRNA levels in Nup133-/- compared to WT
at day 0: Nuggc
at day 3: Nup210L
Lhx1
D Candidates with lower mRNA levels in Nup133 -/- relative to WT (HM1) at day 3:
Magohb
Wfkkn1
Acta2
E Candidate genes with lower mRNA levels in Nup133 -/- compared to WT (HM1) at day 0:
Lhx1
Apln
Cyp26a1
Igfbp3
Rasgrp1
T

**Figure S1, relative to Figure 1: DEGs analysis in differentiating *Nup133*$^{-/-}$ cells.**

**A.** Volcano plot of the RNA-seq analysis carried out in cells at day 2 of neuroectodermal differentiation, showing differentially expressed genes (DEGs), by fold change (log2FC of *Nup133*$^{-/-}$ compared to *WT*) and significance (adj. p.Val presented on a -log10 scale). Significantly upregulated DEGs (adj. p-value<0.05, logFC>2) and downregulated DEGs (adj. p-value<0.05, logFC<-2) are represented by red and blue dots, respectively, if their average normalized expression in log2(CPM) is above 1. The other genes are represented as grey dots when their average expression is below 1 and otherwise as black dots. For this time point, the p-value of several genes is higher (less significant) due to the missing sample (see Materials and Methods). The name of DEGs assessed by RT-qPCR are indicated in blue or red. The names of other relevant DEGs are indicated in grey. **B-E.** mRNA levels of the indicated candidate genes (normalized to *TBP*) were measured by RT-qPCR in pluripotent mESCs or at day 3 of differentiation (see text for details). Data are presented as the mean±S.D. Each dot represents an individual experiment.

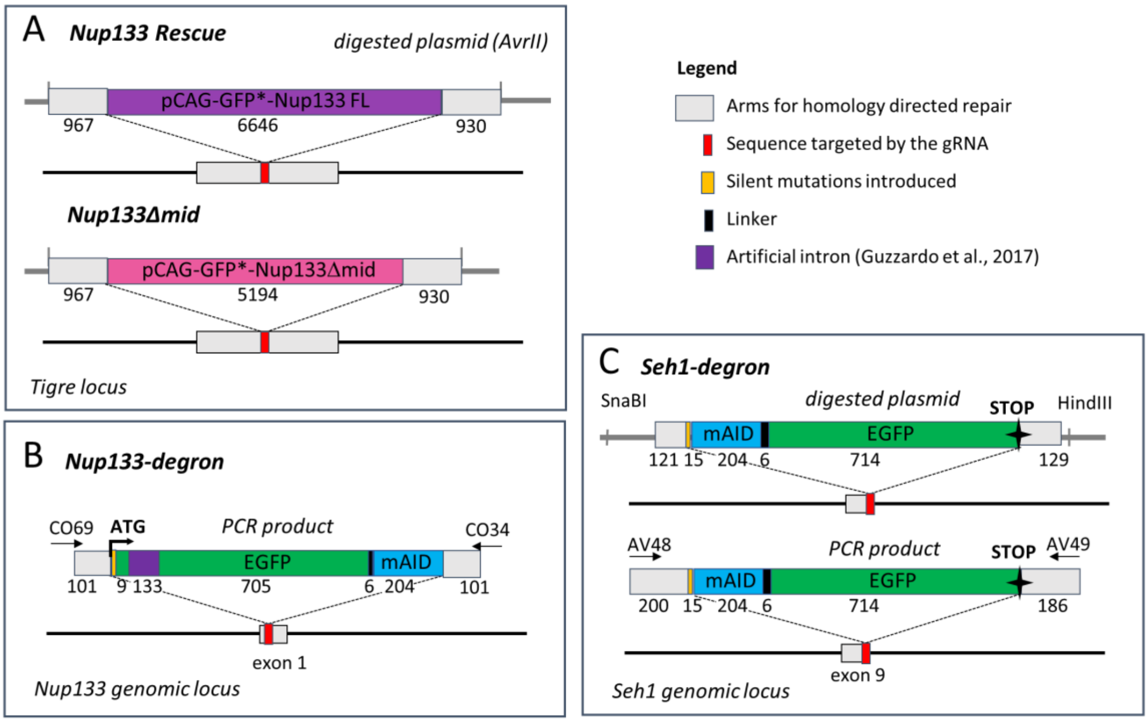

**Figure S2, related to Materials and Methods and Tables S1, S3 and S4: Schematics of CRISPR-Cas9-mediated cell line establishment via homologous recombination.** *Nup133 Rescue* and *Nup133Δmid* (**A**), *Nup133-degron* (**B**), and *Seh1-degron* **(C).** Fragment lengths are indicated in base pairs. mAID: 7kDa mini-auxin Inducible Degron (mAID) sequence (Natsume et al., 2016). Note that two different GFP-coding sequences were used in this study: GFP*, initially described in Harkins et al. (2017), contains, in addition to the S65T and F64L substitutions present in EGFP, the V163A substitution that is expected to confer it a brighter and more stable signal. In addition, the leucine introduced at position 231 during the creation of EGFP (Tsien, 1998) is reverted to the original Histidine.

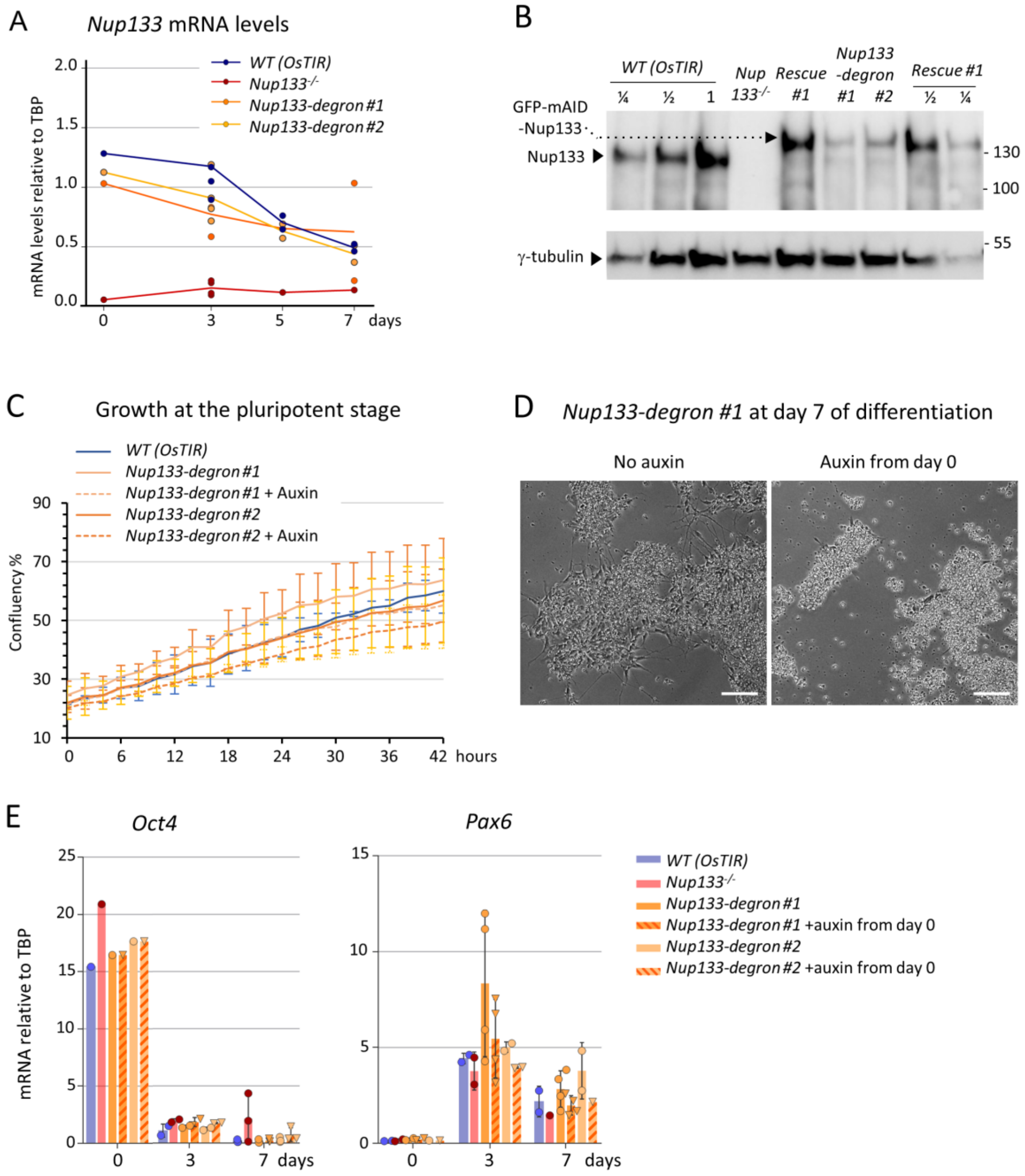
A
Nup133 mRNA levels
mRNA levels relative to TBP
WT (OsTIR)
Nup133-/-
Nup133-degron #1
Nup133-degron #2
days
B
WT (OsTIR)
Nup 133-/-
Rescue #1
Nup133 -degron #1 #2
Rescue #1
GFP-mAID -Nup133
Nup133
γ-tubulin
130
100
55
C
Growth at the pluripotent stage
WT (OsTIR)
Nup133-degron #1
Nup133-degron #1 + Auxin
Nup133-degron #2
Nup133-degron #2 + Auxin
Confluency %
hours
D
Nup133-degron #1 at day 7 of differentiation
No auxin
Auxin from day 0
E
Oct4
Pax6
mRNA relative to TBP
WT (OsTIR)
Nup133-/-
Nup133-degron #1
Nup133-degron #1 +auxin from day 0
Nup133-degron #2
Nup133-degron #2 +auxin from day 0
days

**Figure S3, relative to Figure 4: Characterization of the *Nup133-degron* cell lines**.
**A.** mRNA levels of *Nup133* during differentiation, quantified by RT-qPCR and normalized to *TBP* level, are comparable in the *WT (OsTIR)* and *Nup133-degron* cell lines. **B.** Western blot showing the different expression levels of Nup133 in the *OsTIR, Nup133-degron* and *Rescue* neuronal progenitors at day 5. γ-tubulin was used as loading control. 1/2 and 1/4 dilutions of the *WT (OsTIR)* and *Rescue* extracts were also loaded. **C.** Growth of *Nup133-degron* is comparable to *WT* and not altered by addition of auxin (added at t=0) in pluripotent mESCs. Confluence values were obtained using the IncuCyte® live cell imager software (Essen Biosciences, Ann Arbor, MI) as previously described (Gonzalez-Estevez, Verrico et al., 2021). Error bars correspond to standard deviations from 2 independent experiments. **D.** Images of *Nup133-degron* cells, treated or not with auxin from day 0, were acquired at day 7 of differentiation using a widefield microscope. Neuronal rosettes and axons are visible in untreated cells while major cell death is observed in auxin-treated *Nup133-degron* cells. Scale bars, 100μm. **E.** RT-qPCR analyses show that auxin-treated *Nup133-degron* cells properly repress the pluripotency marker *Oct4* and induce the neuronal progenitor marker *Pax6* when induced to differentiate towards neuroectoderm.

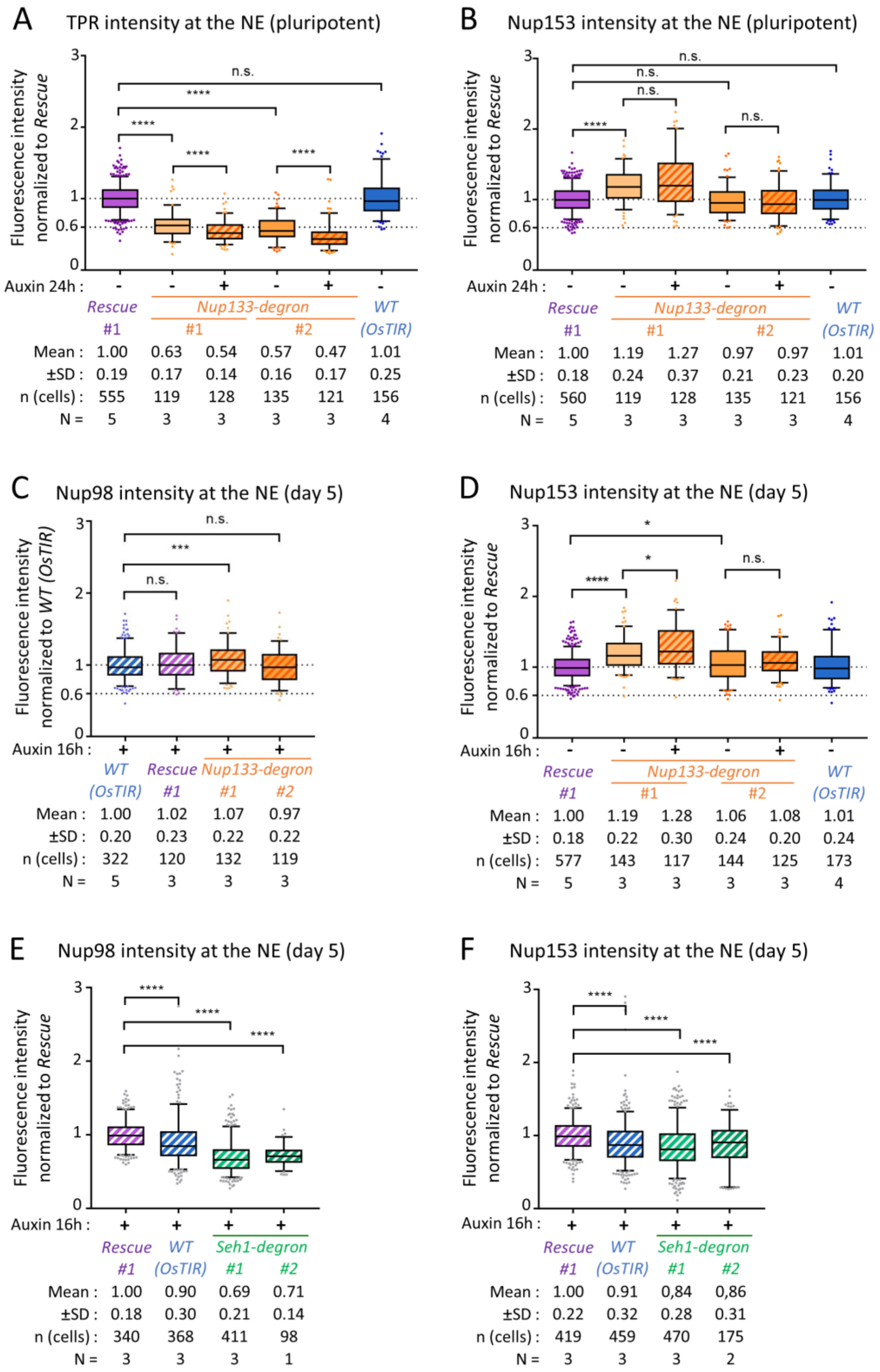
A
TPR intensity at the NE (pluripotent)
Fluorescence intensity normalized to Rescue
n.s.
****
Auxin 24h : - - + - + -
Rescue #1
Nup133-degron #1 #2
WT (OsTIR)
Mean : 1.00 0.63 0.54 0.57 0.47 1.01
±SD : 0.19 0.17 0.14 0.16 0.17 0.25
n (cells) : 555 119 128 135 121 156
N = 5 3 3 3 3 4
B
Nup153 intensity at the NE (pluripotent)
Fluorescence intensity normalized to Rescue
n.s.
****
Auxin 24h : - - + - + -
Rescue #1
Nup133-degron #1 #2
WT (OsTIR)
Mean : 1.00 1.19 1.27 0.97 0.97 1.01
±SD : 0.18 0.24 0.37 0.21 0.23 0.20
n (cells) : 560 119 128 135 121 156
N = 5 3 3 3 3 4
C
Nup98 intensity at the NE (day 5)
Fluorescence intensity normalized to WT (OsTIR)
n.s.
***
Auxin 16h : + + + +
WT (OsTIR)
Rescue #1
Nup133-degron #1 #2
Mean : 1.00 1.02 1.07 0.97
±SD : 0.20 0.23 0.22 0.22
n (cells) : 322 120 132 119
N = 5 3 3 3
D
Nup153 intensity at the NE (day 5)
Fluorescence intensity normalized to Rescue
*
****
n.s.
Auxin 16h : - - + - + -
Rescue #1
Nup133-degron #1 #2
WT (OsTIR)
Mean : 1.00 1.19 1.28 1.06 1.08 1.01
±SD : 0.18 0.22 0.30 0.24 0.20 0.24
n (cells) : 577 143 117 144 125 173
N = 5 3 3 3 3 4
E
Nup98 intensity at the NE (day 5)
Fluorescence intensity normalized to Rescue
****
Auxin 16h : + + + +
Rescue #1
WT (OsTIR)
Seh1-degron #1 #2
Mean : 1.00 0.90 0.69 0.71
±SD : 0.18 0.30 0.21 0.14
n (cells) : 340 368 411 98
N = 3 3 3 1
F
Nup153 intensity at the NE (day 5)
Fluorescence intensity normalized to Rescue
****
Auxin 16h : + + + +
Rescue #1
WT (OsTIR)
Seh1-degron #1 #2
Mean : 1.00 0.91 0,84 0,86
±SD : 0.22 0.32 0.28 0.31
n (cells) : 419 459 470 175
N = 3 3 3 2

**Figure S4, related to Figure 3, B-D and Figure 4D: Nucleoporin densities at the nuclear envelope in *Nup133-degron* and *Seh1-degron* cell lines.**

Fluorescence intensity at the nuclear envelope of Tpr (**A**), Nup153 (**B, D, F**) and Nup98 (**C, E**) was quantified in *Nup133-degron* cells at the pluripotent stage **(A, B),** in *Nup133-degron* cells at day 5 of differentiation (**C, D**), and in *Seh1-degron* cells at day 5 of differentiation (**E, F**) and is presented as box-plots. Cells were treated with auxin (+) or ethanol (-) as control for 24 h or 16h as indicated. Values were normalized in each field to the *Nup133-Rescue* (**A, B, D, E, F**) or *WT (OsTIR*) cells (**C**). ****: p-value<0.0001; ***: p-value<0.001; **: p-value<0.01; *: p-value<0.05; n.s.: non-significant in Mann-Whitney test. Note that Nup153 levels at the NE are mildly affected in a similar manner in pluripotent and differentiated *Nup133-degron* cell lines, with a mild increase observed mainly in *Nup133-degron #1* cells treated or not with auxin (compare panels B and D). In contrast, Tpr levels are strongly decreased in both cell lines (compare panel B with Figure 1E). Note also that in auxin-treated *Seh1-degron* cells, the decreased NE level of Nup98 (panel **E**) (and of Tpr, see Figure 3D) but not of Nup153 (panel F) is consistent with a decreased NPC density combined with an increased stoichiometry of Nup153 per NPC.

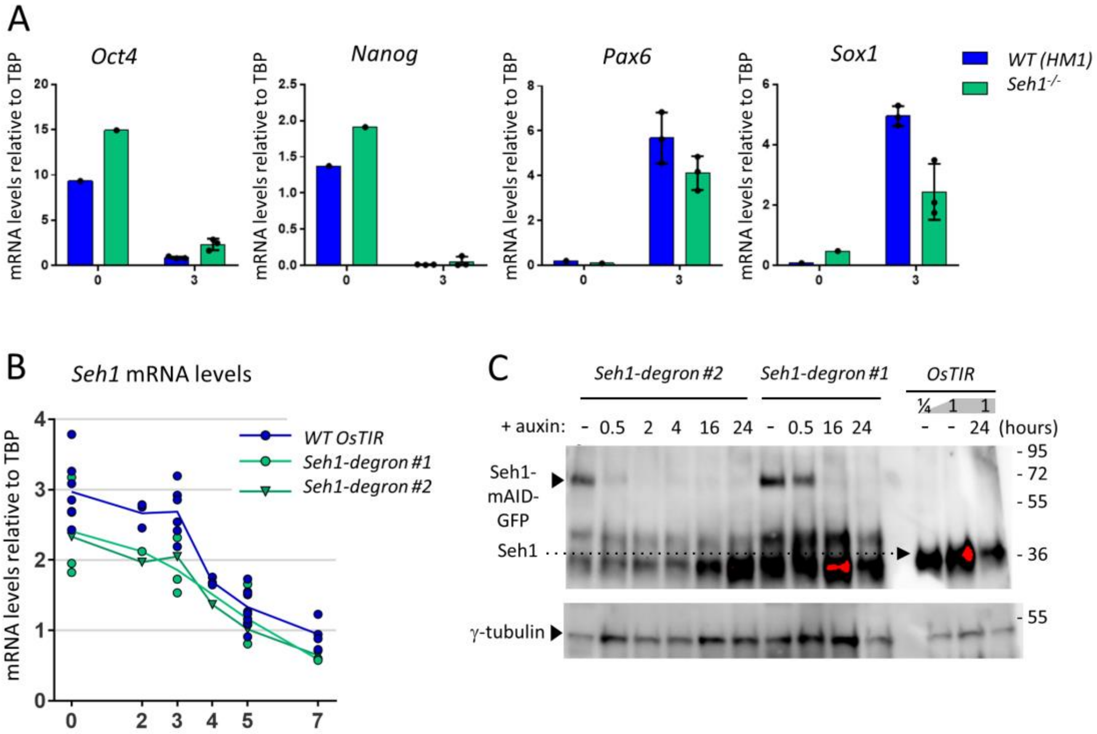

**Figure S5, relative to Figure 5: Characterization of the *Seh1$^{-/-}$* *and Seh1-degron* cell lines**
**A.** mRNA levels of pluripotency (*Oct4*, *Nanog*) and neuronal progenitor (*Pax6*, *Sox1*) markers were analyzed by RT-qPCR in *WT (HM1)* and *Seh1$^{-/-}$* mESCs at the pluripotent stage (n=1) and after 3 days of neuroectodermal differentiation (n=3). **B.** *Seh1* mRNA levels during differentiation, quantified by RT-qPCR and normalized to *TBP* level, are comparable in the *WT (OsTIR)* and the two *Seh1-degron* cell lines (#1 and #2). **C.** Western blot showing Seh1 levels upon auxin treatment in cells at day 5 of differentiation. The membrane was first hybridized with the Seh1 antibody, then stripped to perform γ-tubulin hybridization.

**Table S1, related to Experimental Procedures: Cell lines used in this study**

| NAME | **SOURCE** (IF PUBLISHED OR COMMERCIAL) **OR DESIGN FOR CELL LINES GENERATED IN THIS STUDY** | IDENTIFIER | Plate ref. | MUTATIONS (genomic sequences set at 1 for the first ATG) | Chromosome count/ Karyotype |
|---|---|---|---|---|---|
| **MEF (DR4)** | Applied StemCell | ASF-1001 | - | - | - |
| ***WT* (1A4)** | LACY lab (Souquet et al, 2018) | #1A4 | - | *$Sox1^{gfp/+}$* | 40 Chr (68%)<br>(XY) |
| ***merm #319*** | LACY lab (Souquet et al, 2018) | #319 | - | *$Nup133^{merm/merm}$; $Sox1^{gfp/+}$* | 42 Chr<br>XX, +8, +12 |
| ***WT* mESCs (HM1)** | ThermoFisher Scientific (Selfridge et al., 1992) | MES4303 | - | - | 40 Chr<br>(XY) |
| ***$Nup133^{-/-}$*** | DOYE lab - Generated by CRISPR/Cas9 editing.<br>#14 was described in (Souquet et al., 2018);<br>#19 is a distinct clone that arose from the same editing<br>Parental cell line: HM1<br>Cas9 & gRNA plasmids:<br>pX-102: (two tru gRNAs flanking Nup133 exon 2 andCas9-mCherry)<br>AND pX-98: two gRNAs within mNup133 exon 6 and nCas9-EGFP | *$Nup133^{-/-}$*<br>(#14) | #14 | Allele 1: 35 bp deletion in exon 6 (Δ11094-111128), leading to a frameshift after aa 259.<br>Allele 2: large deletion encompassing exons 2-6 (Δ2525-11143) causing a frameshift after aa 59.<br>Very low mRNA levels due to premature stop codons. No protein detected by western blot. | 40 Chr |
| | | *$Nup133^{-/-}$*<br>(#19) | #19 | Allele 1: 34 bp deletion in exon 6 (Δ11095-11128), leading to a frameshift after aa 260.<br>Allele 2: deletions removing (i) part of exon 2 including its 5' splice site (Δ2539-2719) and (ii) exon 3- 6 (Δ4075-11094). Expected to cause a frameshift after aa 59.<br>Very low mRNA levels due to premature stop codons. No protein detected by western blot. | ND |
| ***$Seh1^{-/-}$*** | DOYE lab<br>(Gonzalez-Estevez, Verrico *et al*, 2021) | *$Seh1^{-/-}$* #1 | A18 | As described in (Gonzalez-Estevez, Verrico *et al*, 2021). | 40 Chr<br>(>60%) |

| | | | | | |
|---|---|---|---|---|---|
| ***Nup133 Rescue*** | Parental cell line: *$Nup133^{-/-}$* (#14)<br>Cas9 & gRNA plasmid: pU6-sgTIGRE_CBh-Cas9-T2A-mCherry-3UTR (#2061)<br>HR template: linearized (with AvrII) pTIGRE-CAG-GFP*mNup133FL (#2103) | *Nup133-Rescue #1* | B11 | Insertion of pCAG-GFP*-Nup133 in one of the *Tigre* alleles | ND |
| | | *Nup133-Rescue #2* | F5 | Insertion of pCAG-GFP*-Nup133 in one of the *Tigre* alleles | 41 Chr (40%)<br>40 Chr (33%)<br>42 Chr (23%) |
| | | *Nup133-Rescue #3* | G5 | Insertion of pCAG-GFP*-Nup133 in one of the *Tigre* alleles | ND |

| | | | | | |
|---|---|---|---|---|---|
| ***Nup133-Δmid*** | Parental cell line: *Nup133*$^{-/-}$ (#14)<br>Cas9 & gRNA plasmid: pU6-sgTIGRE_CBh-Cas9-T2A-mCherry-3UTR (#2061)<br>HR template: linearized (with AvrII) pTIGRE-CAG-GFP*mNup133dMid (#2102) | *Nup133-Δmid #1* | D3 | Insertion of pCAG-GFP*-Nup133Dmid in one of the *Tigre* alleles | 40 Chr (71%) |
| | | *Nup133-Δmid #2* | F4 | Insertion of pCAG-GFP*-Nup133Dmid in one of the *Tigre* alleles | 40 Chr (70%) |
| | | *Nup133-Δmid #3* | C5 | Insertion of pCAG-GFP*-Nup133Dmid in one of the *Tigre* alleles | 40 (73%) |

| | | | | | |
|---|---|---|---|---|---|
| ***WT (OsTIR)*** | As described in (Gonzalez-Estevez, Verrico *et al*, 2021), but a distinct clone<br>Parental cell line: HM1<br>Cas9 & gRNA plasmid: pU6-sgTIGRE_CBh-Cas9-T2A-mCherry-3UTR (#2061)<br>HR template: linearized (with EcoRV) TIGRE HR-pCAG-OsTir-T2A-NeoR-TIGRE HR (#2064) | *OsTIR* | 5F | Insertion of pCAG-OsTir-T2A-NeoR in one of the *Tigre* alleles. | 40 Chr |

| | | | | | |
|---|---|---|---|---|---|
| ***Nup133-degron (GFP-mAID-Nup133)*** | Parental cell line: *OsTIR (5F)*<br>Cas9 & gRNA plasmid: Cas9HF-Cherry-1gRNA-degron-Nup133<br>HR template: PCR product: HR-EGFP-mAID-mNup133 (gRNA resistant)-HR; amplification of plasmid #2108 with primers CO69-CO34 | *Nup133-degron #1* | B7 | Allele 1: insertion of EGFP-mAID after the ATG of Nup133<br>Allele 2: 10bp deletion at +1 | 40 (100%) |
| | | *Nup133-degron #2* | C8 | Allele 1: insertion of EGFP-mAID after the ATG of Nup133<br>Allele 2: 1bp insertion at +10 | 40 (74%) |

| | | | | | |
|---|---|---|---|---|---|
| ***Seh1-degron (Seh1-mAID-GFP)*** | Parental cell line: *OsTIR (5F)*<br>Cas9 & gRNA plasmid: **#2115_**Cas9mCherry_gRNA-Seh1Cterm<br>gRNA sequence: AGCTGAGTACAAGCTAGC<br>HR template: PCR product: amplification of plasmid **#2120** [Seh1 intron8/exon9 HR-mAID-EGFP-mSeh1 exon9 UTR (gRNA resistant)-HR] with primers AV48/AV49 (for clone D2) OR **#2120** digested with SnaBI and HindII (clone E9) | *Seh1-mAID-GFP* #1 | E9 | Alleles 1 and 2: Seh1–mAID-EGFP<br>(only one band detected and sequenced, corresponding to the AID predicted allele).<br>Note that both alleles carry a 207 bp endoduplication in intron 8, 28 bp before exon 9 that however did not affect Seh1 expression compared to *Seh1-mAID-GFP* #2 cells | 40 Chr (>86%) |
| | | *Seh1-mAID-GFP* #2 | D2 | Only one band detected by PCR<br>Seh1–mAID-EGFP as predicted | 40 Chr (>76%) |

| **Table S2. Plasmids used in this study** | | |
|---|---|---|
| **Plasmid** | **Source** | **Identifier (in lab)** |
| pU6_CBh-Cas9-T2A-mCherry Tigre 3UTR | From P. Navarro Gil and N. Festuccia (Festuccia et al., 2019) | #2061 |
| pTIGRE-CAG-GFP*mNup133dMid | This paper | #2102 |
| pTIGRE-CAG-GFP*mNup133FL | This paper | #2103 |
| Cas9HF-Cherry-1gRNA-degron-Nup133 | This paper | #2109 |
| CMV-EGFP-IntronVariant1-mAID-hSeh1cDNA | This paper | #2108 |
| Cas9HF-Cherry-1gRNA-degron-Seh1Cterm | This paper | #2115 |
| mAID-EGFP-mSeh1-exon9-UTR | This paper | #2120 |
| pCAG-GFP*-miniNup210L (SP-GFP*-TM-Cter) | This paper | #2130 |

Plasmids used in this study were either previously published or generated using standard molecular cloning techniques as previously described in (Gonzalez-Estevez, Verrico et al., 2021). Plasmid maps are available upon request. See details about GFP* in the legend to Figure S2.

| **Table S3. gRNA sequences used in this study** | | |
|---|---|---|
| **Identifier** | **Sequence** | **Source** |
| sg-TIGRE | ACTGCCATAACACCTAACTT | (Gonzalez-Estevez, Verrico et al., 2021) |
| gRNA-degron-Nup133 | GTTCGCGGAGAGGAGACGCT | This paper |
| gRNA-Seh1Cterm | AGCTGAGTACAAGCTAGC | This paper |

| **Table S4. Primers used to generate homology-directed repair templates for *Nup133-degron* and *Seh1-degrons*** | | |
|---|---|---|
| **Name** | **Sequence** | **Source** |
| CO69 (FW, plasmid #2108) | CCTCAGGTGTTCAAGCTCCGGGCGCGCGAGGTTCTCGCTATTAGCCC GCGAGTGCCGCTTCCTCCACGCTCTCTGCAAACATGGTGAGCAAGGT AAGTATC | This paper |
| CO34 (RV, plasmid #2108) | GTGGACGTGGGCCCGATCCCGACTAGCGGACCGCGTCGGGTCCCCG GCCCTGGGGTTCGCGGGCTGGAGACGGACGGAAAGGATCTGAGTCC GGATTTATA | This paper |
| AV48 (FW, #2120 plasmid) | GAACAAATTATTTATGAAGGAAAGCATAGTAGAGTTAATTTTTAAA AATCTGGTTTTAGGTTACAGCTTTAGTTCTGTGTTATTTTCTTCCATAA ATAGC | This paper |
| AV49 (RV, #2120 plasmid) | GGCTGACTAGTACAAAAGTTATATACACTGTCACTTTAAAGG CCTTTTGGACTGTGTGTAGTAGTGAACGGGCTGCTGGTGCATCT TGGAAGAGATCAAAC | This paper |

| **Table S5: qPCR primers used in this study** | | | |
|---|---|---|---|
| **Name** | **forward** | **reverse** | **Identifier (in lab) #** |
| TBP | GAAGAACAATCCAGACTAGCAGCA | CCTTATAGGGAACTTCACATCACAG | 11/12 |
| Nup133-Nter (exons 4-6) | GATTTGGTGGCCCTGTCTTA | GAAACTTCCTCCCTGCACTG | 3/4 |
| Nup133-Mid (exons 13-14) | GACAAGGCCGTGACTCAGAT | CAAGCCGACTTGGTGAAGGA | 330/331 |
| Oct4 | TGCCCAGCATCACTATTTCA | GAAGCGACAGATGGTGGTCT | 17/18 |
| Nanog | TTGCTTACAAGGGTCTGCTACT | ACTGGTAGAAGAATCAGGGCT | 43/44 |
| Sox1 | TGGGTCTCAGAAGGAGGATG | TGGGATAAGACCTGGGTGAG | 23/24 |
| Pax6 | GGGAAAGACTAGCAGCCAAA | TGAAGCTGCTGCTGATAGGA | 336/337 |
| Lhx1 | CTTTGCAGCTACACCCAAGC | CTTGGAGCGTCGATTCTGGA | 440/441 |
| Acta2 | GAGAAGCCCAGCCAGTCG | CTCTTGCTCTGGGCTTCA | 168/168 |
| Nup210L | AGCACTCAATGCTCACGACA | GTTGATGCCAGCACAGCAAA | 215/216 |
| Nup153 | AAGAAAGCCGACAGTGAGGA | TTTGCTGCACCTTGATCAGT | 5/6 |
| Ddx3y | CAATTTTGATTTGCCAAGCG | TTTGTGATGTTCAAATTCCTCTCA | 549/550 |
| Eif2s3y | ATTTGGTGAAAGAAAGCCAGG | GGAGCTCCTTCGGCTACTG | 545/546 |
| Wfikkn1 | GGAAGCCATCCTGGCATGT | CGCACAGTCCTGGTCCC | 434/435 |
| Cyp26a1 | AAGCTCTGGGACCTGTACTGT | CTCCGCTGAAGCACCATCT | 561/562 |
| Igfbp3 | TCAATGTGCTGAGTCCCAGA | TGTCCACACACCAGCAGAAG | 555/556 |
| Nuggc | CATTCCAGGCACAGGAGACT | TCATGGGTCTTTCCTCCAGA | 553/554 |
| T (Brachyury) | CTGGGAGCTCAGTTCTTTCG | GTCCACGAGGCTATGAGGAG | 67/68 |
| Magohb | CCGGACGGGAAGCTTAGATA | AGAGCGTCGTCCTCTTTTGT | 253/254 |
| Alpn | CTCTGGCTCTCCTTGACTGC | GCGCATGCTTCCTTCTTCTA | 293/294 |
| Rasgrp1 | ACCGGATCATCATCTCCTCA | AATTCTTTTCCAGGGCATCC | 297/298 |

| Table S6: Antibodies used in this study | | | | |
|---|---|---|---|---|
| **Primary Antibodies** | **Usage** | **Concentration** | **reference** | **source** |
| Mouse monoclonal antibody anti-Nup153 (SA1) | IF | 1/5 | #11 | From B. Burke (Bodoor et al., 1999) |
| Rabbit polyclonal antibody anti-Tpr | IF | 1/200 | #158 | Abcam (ab84516) |
| Rabbit monoclonal anti Nup98 C39A3 | IF | 1/20 | # | Cell signaling (#2598) |
| Rat monoclonal anti-mouse Nup133 antibody (clone 9C2H8) | IF | 1/100 | #74 | Doye lab (Berto et al., 2018) |
| Rat monoclonal anti-mouse Nup133 antibody (clone 3C11G6) | WB | 1/10 | #75 | Doye lab (Souquet et al., 2018) |
| Rabbit monoclonal antibody anti-Nup133 (EPR10809) | WB | 1/500 | #295 | Abcam (ab181355) |
| Rabbit polyclonal antibody anti-Seh1 | WB | 1/1000 | # | Abcam (ab218531) |
| Mouse anti-gamma-tubulin | WB | 1/2500 | #3 | Abcam (ab11316) |
| **Secondary antibodies** | | | | |
| Cy™5 AffiniPure Donkey anti-mouse | IF | 1/500 | #504 | Jackson ImmunoRes. 715-165-151 |
| Cy™5 AffiniPure Donkey anti-rat | IF | 1/500 | #561 | Jackson ImmunoRes. 712-035-152 |
| Cy™3 AffiniPure Goat anti-rabbit | IF | 1/500 | #519 | Jackson ImmunoRes. 111-165-144 |
| Peroxidase AffiniPure Donkey anti-rabbit | WB | 1/1500 | #499 | Jackson ImmunoRes. 711-035-152 |
| Peroxidase AffiniPure Goat anti-mouse | WB | 1/5000 | #515 | Jackson ImmunoRes. 115-035-068 |
| Peroxidase AffiniPure Goat anti-ratCy™5 AffiniPure Donkey Anti-Rat | WB | 1/10000 | #523 | Jackson ImmunoRes. 112-035-167 |

**References cited in Supplemental Materials and Methods:**

Berto, A., Yu, J., Morchoisne-Bolhy, S., Bertipaglia, C., Vallee, R., Dumont, J., Ochsenbein, F., Guerois, R., Doye, V. (2018). Disentangling the molecular determinants for Cenp-F localization to nuclear pores and kinetochores. EMBO Reports. 19, e44742.

Bodoor, K., Shaikh, S., Salina, D., Raharjo, W. H., Bastos, R., Lohka, M., Burke, B. (1999). Sequential recruitment of NPC proteins to the nuclear periphery at the end of mitosis. J. Cell Sci. 112, 2253-2264.

Festuccia, N., Owens, N., Papadopoulou, T., Gonzalez, I., Tachtsidi, A., Vandoermel-Pournin, S., Gallego, E., Gutierrez, N., Dubois, A., Cohen-Tannoudji, M., Navarro, P. (2019). Transcription factor activity and nucleosome organization in mitosis. Genome Research. 29, 250-260.

Gonzalez-Estevez, A., Verrico, A., Orniacki, C., Reina-San-Martin, B., and Doye, V. (2021). Integrity of the short arm of the nuclear pore Y-complex is required for mouse embryonic stem cell growth and differentiation. J. Cell Sci. 134, jcs258340.

Harkins, H.A., Page, N., Schenkman, L.R., De Virgilio, C. Shaw,S., Bussey, H.,and Pringle, J.R. (2001) Bud8p and Bud9p, proteins that may mark the sites for bipolar budding in yeast. Mol. Biol. Cell 12,2497-518.

Guzzardo, P. M., Rashkova, C., Dos Santos, R. L., Tehrani, R., Collin, P., Bürckstümmer, T. (2017). A small cassette enables conditional gene inactivation by CRISPR/Cas9. Scientific Reports. 7, 16770.

Natsume, T., Kiyomitsu, T., Saga, Y., Kanemaki, M. (2016) Rapid Protein Depletion in Human Cells by Auxin-Inducible Degron Tagging with Short Homology Donors. Cell Reports. 15, 210-218.

Selfridge, J., Pow, A.M., Mcwhir, J., Magin, T. M., Melton, D. W. (1992) Gene Targeting Using a Mouse HPRT Minigene/HPRT-Deficient Embryonic Stem Cell System: Inactivation of the Mouse ERCC-1 Gene. Somatic Cell and Molecular Genetics. 18, 325-336.

Souquet, B., Freed, E., Berto, A., Andric, V., Audugé, N., Reina-San-Martin, B., Lacy, E., Doye, V. (2018) Nup133 Is Required for Proper Nuclear Pore Basket Assembly and Dynamics in Embryonic Stem Cells. Cell Reports. 23, 2443-2454.

Tsien, R.Y. (1998) The green fluorescent protein. Annu. Rev. Biochem. 67, 509-544
